# Optimal tuning not required: birds communicate effectively with mismatched auditory filters

**DOI:** 10.1101/2025.07.29.667030

**Authors:** Matías I. Muñoz, Wouter Halfwerk

**Affiliations:** Department of Ecological Sciences, Vrije Universiteit Amsterdam, De Boelelaan 1085, Amsterdam, 1081 HV, The Netherlands; Department of Integrative Biology, Oklahoma State University, Stillwater, OK 74078, U.S.A.

**Keywords:** Bird vocal communication, auditory tuning, matched-filter hypothesis, allometric scaling, diffuse coevolution

## Abstract

Birds have one of the most remarkable vocal capacities of vertebrates. Yet we know little about the relationship between their hearing and their vocal capacities at large phylogenetic scales, and whether these traits coevolve in a tight or loose fashion. Here we collected vocal and auditory data for 72 bird species from 15 different orders, and tested if hearing is finely tuned to the spectral composition of their vocalizations (i.e., the matched-filter hypothesis). We found bird audition not to be predictive of vocal frequency, and mismatches between hearing and vocalizations to be widespread. We show that the source of these mismatches is allometric: the auditory tuning and the spectral composition of vocalizations scale with body weight to different degrees. Despite the general frequency mismatches, bird hearing remains remarkably sensitive to the main spectral component of their vocalizations. Our results challenge the view that sensory systems must be finely tuned only to conspecific signals for communication to be effective. Our results suggest that selection for a broad auditory tuning has allowed birds to effectively communicate with conspecifics while at the same time permitting general environmental awareness using auditory cues. We argue that the relationship between hearing and vocalizations in birds is better described by the concept of diffuse coevolution, where organism evolve in relation to their broader ecological context instead than on a strictly dyadic fashion.

**Significance Statement:** Animal communication theory often states that signalers and perceivers coevolve, yet the spread and nature of this relationship remains unclear. We tested whether the auditory tuning of birds is tightly matched to the frequency content of their vocalizations -an expected outcome of strong coevolution and the matched-filter hypothesis. We show that although birds hearing does not precisely align with their vocalizations, birds are still highly sensitive to the sounds they produce. This is due to the broad tuning of the avian auditory systems, which supports effective conspecific communication while also allowing the detection of other ecologically relevant sounds. Thus, birds exemplify diffuse rather than tightly coupled coevolution in communication systems.

## Introduction

In animal communication, signalers and perceivers are often said to coevolve (1–3). But what does this actually mean? In one of the earliest and most remarkable examples of coevolutionary reasoning, Darwin famously predicted the existence of an undescribed moth with an extraordinarily long proboscis based solely on the structure of an orchid with an extraordinarily long nectar tube (4). This classic example raises the question: are signalers and perceivers entangled in a similar coevolutionary dynamic? If so, and by analogy with Darwin’s moth, does this mean that knowledge of an animal’s sensory capacities is sufficient to infer the structure of its signals?

The “matched-filter” hypothesis proposes that a perfect match between the sensory capacities of perceivers and the structure of animal signals would maximize signal-to-noise ratio, therefore facilitating the detection of conspecifics under noisy or in highly attenuating conditions (4). In the acoustic domain, frogs are among the most outstanding examples of matched filters, and their auditory systems are finely tuned to the peak frequency of their advertisement calls (6). Another notable example of auditory-vocal matching are bats, where the peak frequency of their echolocation calls closely matches their high-frequency hearing range, whereas their low-frequency hearing sensitivity matches the frequency of pup isolation calls (7). These comparative studies indicate a general benefit of matched-filters for taxa that rely on vocalizations for reproductive, parental and navigation behaviors.

But tuning a sensory system finely only to conspecific signals is not necessarily optimal, as such close correspondence between signalers and perceivers will necessarily reduce the ability to detect other ecologically relevant stimuli. Many animals rely on their senses to detect predators or parasites, find food or shelter, or gather other relevant environmental cues (8–16). These important functions may be compromised when sensory filters are exclusively matched to conspecific signals. How animals navigate this trade-off between effective signal detection and broader environmental awareness remains however a key question in sensory ecology.

Birds are exceptional vocalizers that rely on vocalizations for parent-offspring communication, mate attraction, territorial interactions and in a few species even for navigation. Although vocal behavior has been a major focus of avian bioacoustics research (17, 18), we know surprisingly little about the degree of matching between avian vocalizations and avian audition at large phylogenetic scales. Only a handful of taxonomically-restricted studies have explored this relationship from a multi-species perspective(19, 20), and these have sometimes reached contrasting conclusions (21, 22). Despite the extensive body of literature on bird vocal communication, there is no general consensus about the relationship between the vocal and auditory capacities of birds, and whether avian signalers and perceivers are entangled in the reciprocal selection dynamics that characterize coevolutionary processes.

To address this gap, we collected from the literature the audiograms and body mass for 72 bird species spanning 15 different orders, and analyzed 108 different types of vocalizations from these species (**Fig. 1A**). With this data we quantified the degree of auditory-vocal matching across birds, and the effect of body weight on this matching. From the hearing curve of each species we measured the (i) best hearing frequency, (ii) the low-frequency hearing limit and (iii) the high-frequency hearing limit. From the vocalizations we measured their (i) peak frequency, as well as the (ii) low- and (iii) high-frequency limits. We, therefore, obtained equivalent frequency measures from both the spectrum of the vocalizations and the auditory curves. First, we tested for the allometric effect of body weight on both auditory and vocal variables (i.e., auditory and vocal allometries). Second, we tested for associations between vocal and auditory variables while accounting for body weight variation. Then, from the auditory and vocal variables we computed three different metrics of auditory-vocal matching (**Fig. 1B** and **1C**, see *Materials and Methods*), and evaluated the effect of body weight on these three metrics. Our analyses were aimed to test if birds adhere or not to the expectation of the matched-filter hypothesis and tight signaler-perceiver coevolution (i.e., an optimal matching between hearing and vocalizations), and the effect of body weight on auditory-vocal matching.

**Figure 1.**
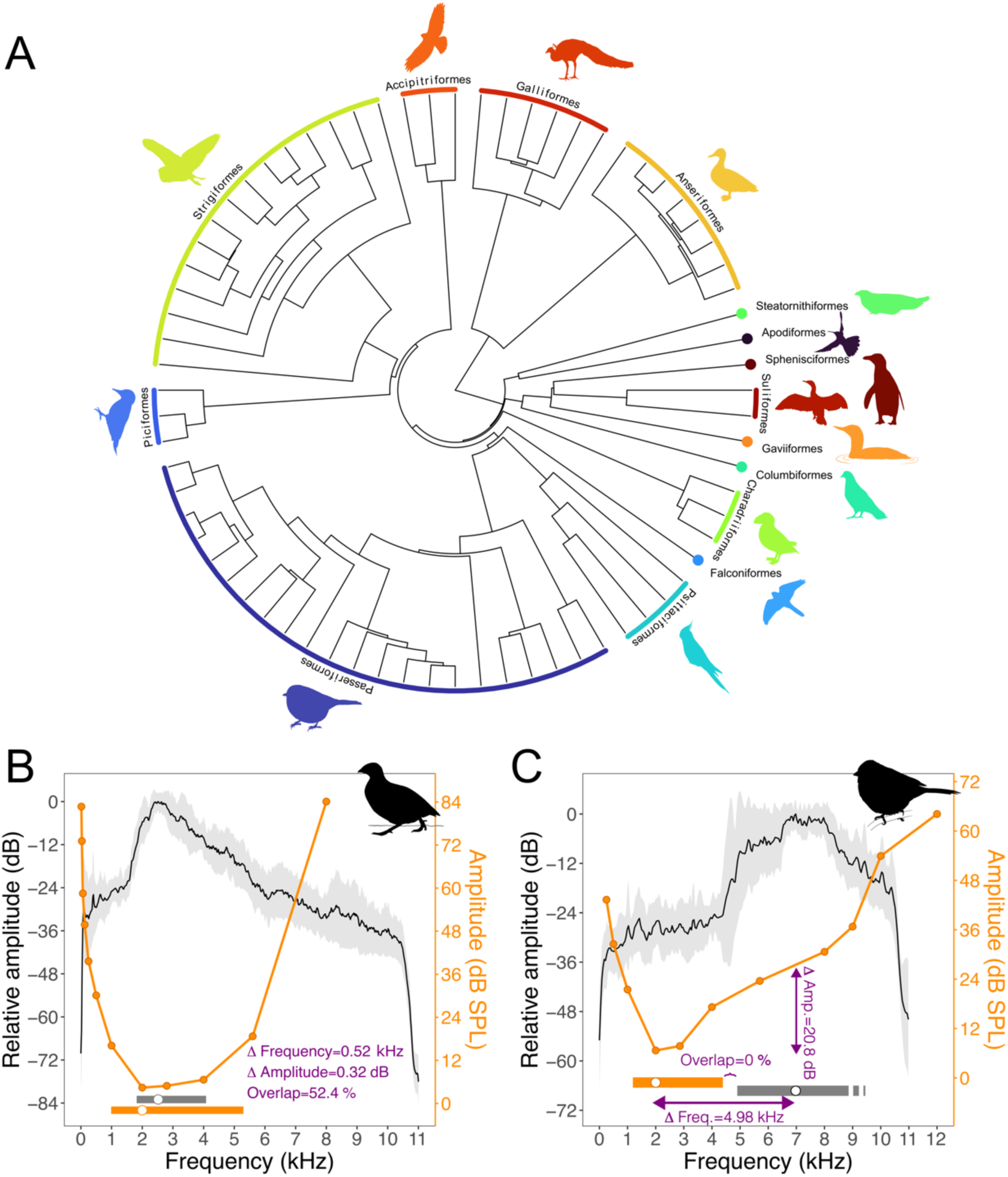
Bird phylogeny and examples of auditory-vocal matching and mismatching. (A) Maximum clade credibility phylogenetic tree of the bird species included in this study. Phylogeny based on 1,000 trees from (65). Example of a species with (B) close auditory-vocal matching (Japanese quail, *Coturnix japonica*) and (C) auditory-vocal mismatch (field sparrow, *Spizella pusilla*). The solid orange points and lines show the audiogram for the species (right axis), and the black line and grey shading show the mean and standard deviation power spectrum of the vocalization (left axis). The horizontal bars at the bottom depict the ranges of hearing (orange) and vocalizations (grey), and the white dots inside of them show the best hearing frequency and peak frequency, respectively. The three metrics of auditory-vocal matching (i.e., Δ frequency, Δ amplitude and bandwidth overlap) are shown inside panels (B) and (C) in violet. ***SI appendix*, Figs. S4 to S18** show equivalent figures for all the birds included in this study.

## Results

### Strong phylogenetic signal in all vocal and most auditory traits

Phylogenetic signal was strong for all vocal variables (averaged to a single value for species with more than one vocalization analyzed): peak frequency (Pagel’s λ = 0.7482, *P* < 0.001), low-frequency (λ = 0.6474, *P* < 0.001), and high-frequency vocal limits (λ = 0.7865, *P* < 0.001). Phylogenetic signal was also strong for the low-frequency (λ = 0.4681, *P* = 0.004) and high-frequency hearing limits (λ = 0.7859, *P* < 0.001), but weak for the best hearing frequency (λ = 0.0001, *P* = 1). We also found strong phylogenetic signal for body weight (λ = 0.6215, *P* < 0.001).

### Auditory and vocal variables scale differently with body weight

First, we tested the allometric scaling of vocal and auditory variables with body weight. We found that both the peak frequency of the vocalizations and best hearing frequency scale negatively with body weight (although only at the 89% credible level for the best hearing frequency), but peak frequency scales more steeply with body weight than best hearing frequency. (**Fig. 2A and B**; ***SI Appendix,* Fig. S1A and B**; vocal slope = -0.53 [95% CI: -0.75, -0.32], *P* < 0.001; auditory slope = -0.16 [89% CI: -0.32, -0.01], *P* = 0.087). A similar difference in the grade of allometric scaling was found for the high-(**Fig.2D and E**; ***SI Appendix*, Fig. S1D and E**) and low-frequency (**Fig. 2G and H**; ***SI Appendix*, Fig. S1G and H**) limits of hearing and vocalizations. These results indicate that an evolutionary change in body weight will have a more pronounced effect on the frequency of vocalizations than on the auditory tuning of birds.

**Figure 2.**
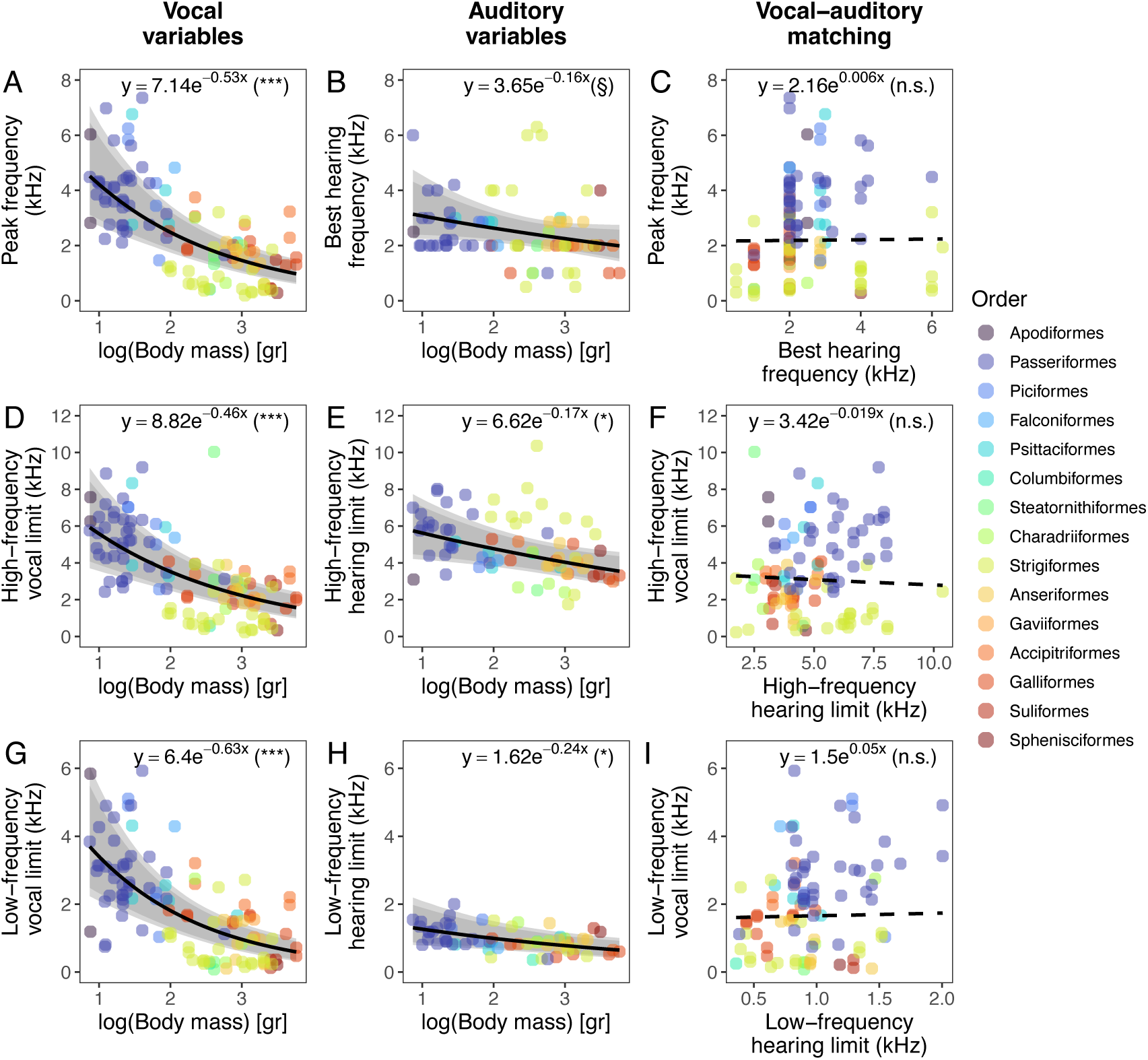
Allometric effects on vocal and auditory variables, and auditory-vocal matching. Effect of body weight on vocal (A, D, G) and auditory variables (B, E, H). Panels (C, F, I) show the absence of relationship between auditory and vocal variables after controlling for variation in body weight. Solid black lines depict significant regression estimates (i.e., with slope 95 or 89% CIs not crossing zero) whereas dashed black lines depict non-significant regressions (i.e., CIs crossing zero, see also ***SI Appendix,* Fig. S1**). The grey shaded area around significant regression lines indicate 89 and 95% credible intervals. The regression equation is shown at the top of each panel, and the significance level (computed from the probability of direction, see *Materials and Methods*) is shown inside brackets (***: *P* < 0.001; *: *P* < 0.05; §: *P* < 0.1; n.s.: *P* > 0.1).

### Audition does not predict the spectral properties of vocalizations

After accounting for variation in body weight (included in the model as a covariate), there was no evidence for an association between best hearing frequency and vocal peak frequency (slope = 0.006 [95% CI: -0.083, 0.093], *P* = 0.889, **Fig. 2C**; ***SI Appendix,* Fig. S1C**). The same was true for the high-frequency hearing and vocal limits (slope = -0.02 [95% CI: -0.10, 0.06], *P* = 0.635, **Fig. 2F**, ***SI Appendix*, Fig. S1IF**), and for the low-frequency hearing and vocal limits (slope = 0.05 [95% CI: -0.35, 0.43], *P* = 0.791, **Fig. 2I, *SI Appendix*, Fig. S1I**). These results indicate that bird auditory curves do not predict the spectral features of their vocalizations.

### Birds have adequate sensitivity to their vocalizations despite frequency mismatches

To quantify auditory-vocal matching, we computed three parameters using frequency and amplitude features from vocalizations and audiograms: (i) Δ frequency—difference between vocal peak frequency and best hearing frequency; (ii) Δ amplitude—difference between hearing sensitivity at the peak vocal frequency and at the best hearing frequency; and (iii) bandwidth overlap—percentage overlap between vocal and hearing frequency ranges (**Fig. 1B, C**; see *Materials and Methods* section).

Across all species, birds showed a close match between the peak frequency of their vocalizations and their best hearing frequency, with a mean Δ frequency (estimated from an intercept-only phylogenetic mixed-effects model) of –0.14 kHz [95% CI: –1.39, 1.15] (**Fig. 3A**). However, orders varied: Anseriformes, Strigiformes, and Suliformes had predominantly negative values, while Accipitriformes, Passeriformes, and Galliformes showed positive values. Therefore, the near zero Δ frequency value we found across all the birds in our sample resulted from different orders having either predominantly positive or negative values, and not from birds necessarily having a tight auditory-vocal matching. For Δ amplitude, birds were on average 5.91 dB [95% CI: 2.62, 13.32] less sensitive at the vocal peak frequency than at their best hearing frequency (**Fig. 3B**), indicating about a 1.97-fold difference in linear sensitivity. Bandwidth overlap between vocal and hearing ranges averaged 30.04% [95% CI: 20.27, 41.24] (**Fig. 3C**), with vocal bandwidths generally narrower than auditory bandwidths across most orders (***SI Appendix*, Fig. S2**). This means that the vocal frequency range of birds typically overlaps with approximately a third of their most sensitive hearing region, although again, there was considerable variation between orders.

**Figure 3.**
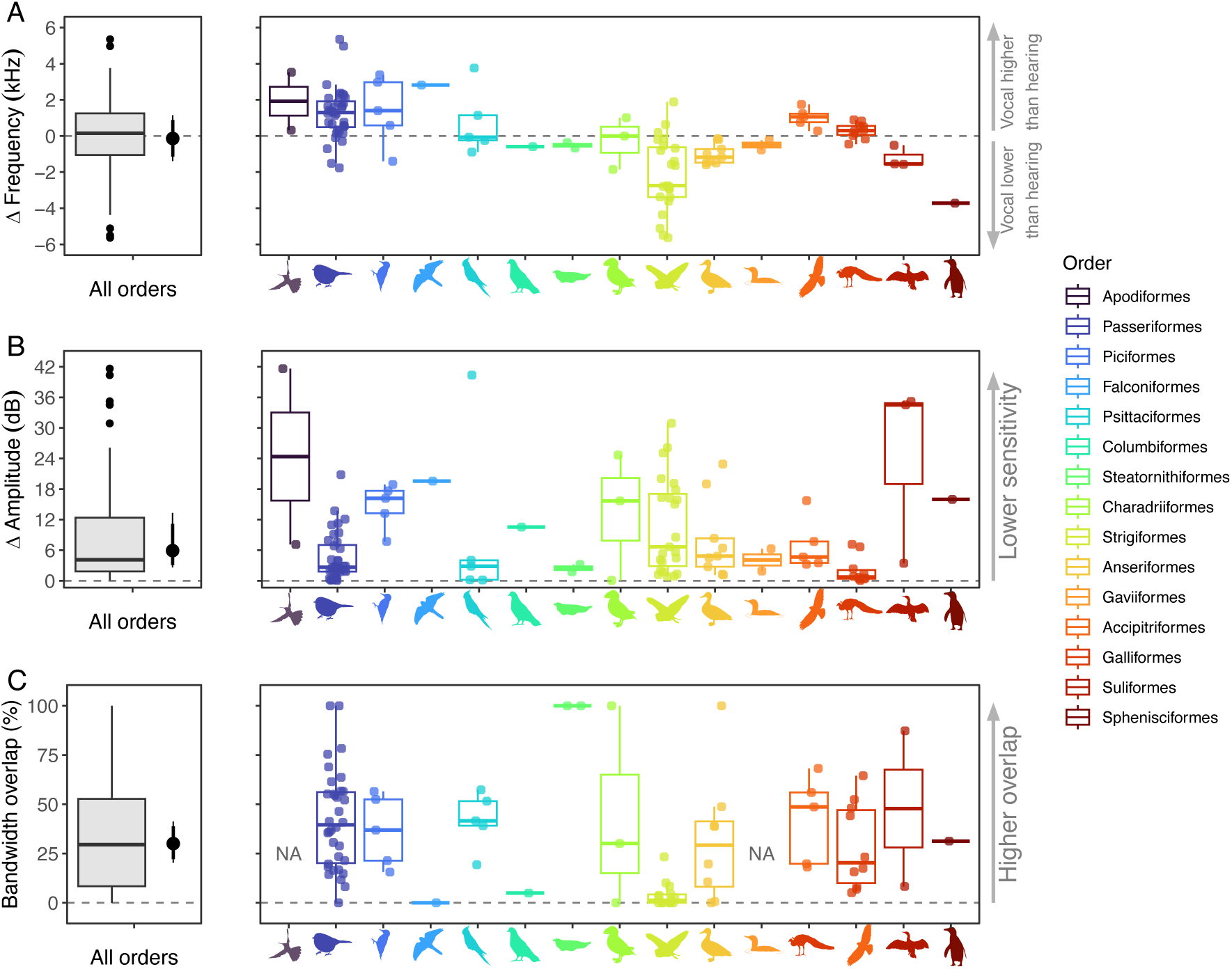
Overview of auditory-vocal matching across birds. (A) Δ frequency (Hz) computed from subtracting the peak frequency of the vocalizations from the best hearing frequency of the audiogram. (B) Δ amplitude (dB) computed from subtracting threshold estimated at the peak frequency from the best hearing threshold. (C) Bandwidth overlap (%) computed from the overlap between the auditory and vocal bandwidths (at 12 dB from the peaks). The left panel shows the data across all bird species in the sample (grey boxplot) and the black point and error bars depict the average posterior estimate from an intercept-only phylogenetic mixed-effects model, and the 89 and 95% credible intervals. The right panel shows the different orders separately, and ordered along the x axis based on their average body weight from lighter (left, Apodiformes) to heavier (right, Sphenisciformes).

### Frequency mismatches depend on body weight

We evaluated if Δ frequency, Δ amplitude and bandwidth overlap scaled allometrically with body weight. We found evidence of a negative association between Δ frequency and body weight (slope = -0.71 [95% CI: -1.40, -0.005], *P* = 0.049, **Fig. 4A, *SI Appendix*. Fig. S3A**), but no effect of body weight on Δ amplitude (slope = -0.10, [95% CI: -0.63, 0.43], *P* = 0.714, **Fig. 4B, *SI Appendix*. Fig. S3B**) or bandwidth overlap (logit slope = -0.02, [95% CI: -0.42, 0.43], *P* = 0.902, **Fig. 4C, *SI Appendix*. Fig. S3C**). This indicates that smaller birds tend to vocalize with peak frequencies higher than their best hearing frequencies (i.e., positive Δ frequencies), while larger birds vocalize with peak frequencies lower than their best hearing frequencies (i.e., negative Δ frequencies).

**Figure 4.**
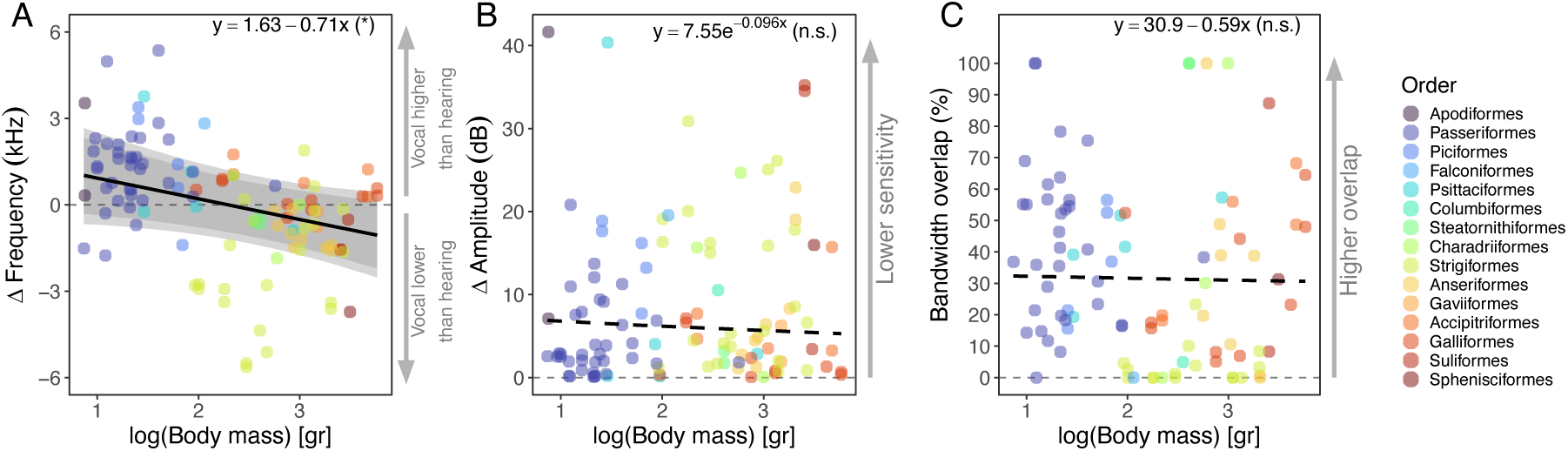
Allometric effects on auditory-vocal matching metrics. Allometric scaling of A) Δ frequency B) Δ amplitude and C) bandwidth overlap with body weight. The dashed lines in panels B) and C) indicate that the model estimate of the slope was not significant at the 95% or 89% credible interval level. The regression equation is shown on the top of each panel, and the significance level (computed from the probability of direction, see *Materials and Methods*) is shown inside brackets (*: *P* < 0.05; n.s.: *P* > 0.10).

## Discussion

Birds have remarkable vocal capacities, yet the degree of matching between their vocal and auditory capacities at broad phylogenetic scales remains poorly understood. Here we show that the tuning of the avian auditory system is not tightly related to the spectral attributes of their vocalizations. Most vocalizations (75%) had peak frequencies within ±2 kHz of the species’ best hearing frequency, and smaller birds tended to vocalize at higher peak frequencies than their best hearing frequency, whereas larger birds vocalized at lower frequencies than their best hearing. Despite these frequency mismatches, birds are still well able to hear their own vocalizations, indicated by our finding that the majority of Δ amplitude values were within 0 and 6 dB (56.5% of analyzed sounds). Also, auditory bandwidths were broader than vocal bandwidths for most avian orders (with the notable exception of the oilbird, who makes broadband noisy shrieks) and around a third (30.6%) of the vocal range overlapped with the auditory range. Our results show that the broadband properties of bird audition allow them to detect a wide range of frequencies without substantially sacrificing auditory sensitivity.

In theory, a close match between perceptual and signaling traits should maximize signal detection, which would be especially advantageous in noisy or highly attenuating environments (5, 23). Comparative studies have confirmed such near-optimal coupling between vocalizations and auditory sensitivity in frogs (6) and bats (7). In frogs, this coupling is so strong that it has been demonstrated not only between signalers and perceivers across populations (24) but also reciprocally between males and females (25)(but see (26, 27)). Bats, as auditory specialists that rely on echolocation for navigation and foraging, exhibit a strong correlation between the peak frequency of their echolocation calls and the frequency of best hearing sensitivity (7). Our finding that the peak frequency of birds is unrelated to their best hearing frequency is in marked contrast to the patterns observed in frogs and bats, and indicates that birds do not adhere to the optimality criteria proposed by the matched-filter hypothesis, at least not for their acoustic signals.

The fact that birds do not adhere to the predictions of the matched-filter hypothesis does not mean that their communication system is suboptimal or deficient. Bird audiograms are typically broader banded than the spectrum of their vocalizations, and therefore frequency mismatches between vocal and auditory peaks did not translate to massive losses of sensitivity. This was confirmed by our finding that Δ amplitude values for most of the vocalizations was within 0 and 6 dB. For reference, a cut-off of 30 dB is typically used to define the highest and lowest limits of avian hearing (28) (in this article we used a cut-off of 12 dB because audio recordings of vocalizations were not always high-quality enough to measure bandwidths at 30 dB, and we wanted to collect equivalent vocal and auditory measures, see *Materials and Methods*). If audiograms and vocal spectra were both narrow banded, then even slight frequency mismatches would lead to drastic decreases in auditory sensitivity (i.e., higher Δ amplitude values), but because this was not the case, birds can detect a wide range of frequencies without sacrificing auditory sensitivity.

### Why do birds hear sounds beyond the range of their own voice?

The broad tuning of the avian auditory system is not only adequate for conspecific communication, but also allows the detection of other ecologically important sounds. We think that selection for broader environmental acoustic cues played an important role in maintaining the broad tuning of avian hearing. For example, owls rely on hearing for prey detection (29), nestlings detect and respond to the sound produced by the footsteps of predators (30), and there is some evidence that albatrosses use infra-sonic geophonies to find the best environmental conditions for flying (12). Furthermore, parent-offspring communication is an important behavior in the life of many birds (31), and birds are well known to rely on the sounds of other species in their surroundings (32, 33). These examples illustrate the diversity of functions for bird hearing beyond adult-adult communication, and help us understand the potential benefits of unmatched filters for ecological situations where animals must do more than simply detect conspecifics.

Birds generally have rich vocal repertoires, larger than the one or two vocalizations we studied here. Great tits (*Parus major*), for example, possess a complex repertoire of vocalizations, some of which they only use during a very short period of time around peak female fertility (34), and some songbirds (Passeriformes) and parrots (Psittaciformes) are well-known to mimic the sounds of other species or even anthropogenic objects (35, 36). Our sampling of vocalizations was limited by the availability of high-quality recordings in the repositories we surveyed, yet we are confident that the vocalizations we analyzed were those most commonly produced by each species. Studying the full repertoire of each species, including the vocalizations of chicks and juveniles, and the rich diversity of non-vocal sounds (e.g., feather flutters, percussions, drumming), will be fundamental to have a complete understanding of the role of hearing in shaping the evolution of acoustic communication in birds. In bats, for example, the low-frequency hearing limit correlates with the peak frequency of pup isolation calls (7), and cochlear elongation in extinct and extant reptiles is correlated with the presence of juvenile vocalizations (37), showing that social communication between parents and off-spring can also play a role in the co-evolution between perceptual and vocal systems.

### Body size constraints signal evolution via sensory biases

The different scaling degrees (i.e., slopes) of vocal and auditory variables with body weight partly explain the mismatches we observed. Our results show that evolutionary changes in body weight lead to a more pronounced shift in vocal frequency than in auditory tuning. One explanation (on a proximate level) for the allometric differences between vocalizations and audition is that their hearing system (i.e., middle and inner ear) is morphologically constrained by its location inside of the skull, whereas the syrinx (i.e., the avian vocal organ) experiences less structural constraints due to its location inside of the chest. Another explanation (on a functional level) could be stabilizing selection on bird’s hearing due to the broad range of environmental sounds they need to detect, which may also explain why their hearing is generally uniform, as most species have similar U-shaped auditory tuning curves with a best hearing frequency between 1 – 4 kHz (18).

Although the causes of the different allometries of vocal and auditory variables are still not clear, their consequences for the evolution of size-dependent mismatches are: bird species at the extreme of body size variation will show more prominent auditory-vocal mismatches. These frequency mismatches create sensory biases that may exert selection on the evolution of acoustic signals simply because some frequencies will be more easily detected than others (38). For relatively small birds, such as most hummingbirds and many passerines, we would, for example, predict intra- and inter-sexual responses to be strongest towards lower frequency songs (*sensu stricto* sensory model of sexual selection) (39). For relatively large birds we would predict sensory biases to favor individuals using higher frequency vocalizations, which may be an opposing force to other selection processes, such as environmental selection favoring lower frequencies in highly attenuating environments (higher frequencies propagate poorly relative to lower frequencies (40), or lower frequencies being a reliable indicator of the (big) size or competitive ability of mates or contenders (41, 42).

## Conclusion

Coming back to our original question: do avian signalers and perceivers coevolve in a similar fashion to the example of Darwin’s moth and orchid? Our results show this is not the case, and bird vocal communication system is not an example of such tight coevolutionary dynamic. Instead, the case of birds is better captured by the concept of diffuse coevolution, in which organisms coevolve with their broader ecological context (e.g., their community) rather than in a strictly dyadic fashion (43, 44). Around 55 years ago, Konishi (21) studied the hearing and vocalizations of ten species of Passeriformes and concluded that “it is obvious that the bird’s ear is not so narrowly tuned to the species song”. We confirm this observation across a larger and more diverse sample of birds. We also add that, despite hearing not being perfectly tuned to their vocalizations, birds are still effective vocal communicators, and broad auditory tuning may allow for greater awareness of diverse sound sources in their environments.

## Materials and Methods

### Hearing data collection

We collected audiogram data from the literature. We mainly searched audiograms obtained through behavioral methods, as these are considered the gold-standard for measuring hearing curves (45). For a number of species, we also collected physiological audiograms obtained either through auditory brain stem responses (ABRs) or cochlear potentials (see ***SI appendix***, **Table S1**). The shape of the audiograms obtained using behavioral and ABRs are generally congruent (46, 47), and the differences between them are mostly related to the absolute amplitude of the thresholds, which are around 20 – 30 dB higher for ABR audiograms. We excluded from our search physiological audiograms obtained at more central levels, like at fore- and mid-brain levels or recordings of single-units (21, 48–50). We also excluded audiograms obtained from juveniles (51). For the species with both behavioral and physiological audiograms available, we prioritized the first over the latter ones. When more than one behavioral audiogram was available for a species, we chose the one that tested a larger number and range of frequencies. Audiograms are generally reported as figures that show the average hearing threshold for each frequency tested for a number of individuals. We extracted the average hearing thresholds in dB SPL (re. 20 μPa) from these figures using the PlotDigitizer web app (version 3.1.5, https://plotdigitizer.com/app). When only median values were available (52), we collected these median values instead of averages. When audiogram data was not averaged in the original articles, we used PlotDigitzer to estimate the thresholds of individual birds and later averaged these individuals to a single audiogram for the species. When auditory thresholds were reported in tables we used these values instead of estimating the thresholds from figures. For 11 species we compared the values reported in tables against the values estimated from the corresponding figures using PlotDigitizer. We found that the thresholds estimated from the figures were accurate, and typically deviated less than 1 dB from the values reported in the tables (average difference: 0.08 dB, SD = 0.730 dB, N = 100). Auditory threshold values were transformed to dB SPL (re. 20 μPa) when these were reported on a different unit or computed based on a different reference (53, 54). For the oilbird (*Steatornis caripensis*), the auditory thresholds reported by Konishi & Knudsen (55) were dB values referenced relative to the best auditory threshold, and therefore values could not be transformed to absolute dB SPL (re. 20 μPa).

### Vocalization data collection

For all the species with audiogram data available, we collected audio-files of their vocalizations mostly from the Xeno-Canto repository (Naturalis Biodiversity Centre, Leiden, The Netherlands). For the oilbird (*Steatornis caripensis*) we collected recordings from Xeno-Canto and the Macaulay library (Cornell Lab of Ornithology, Ithaca, NY, USA). For the Bengalese finch (*Lonchura striata domestica*) recordings were obtained from the supplementary material of Katahira et al., (56). For the zebra finch (*Taeniopygia guttata*) recordings were obtained from the library of vocalizations collected from the colony at the University of California Berkeley (57, 58). Long-tailed finch (*Poephila acuticauda*) recordings were obtained from the Macaulay library.

Most species of birds have rich vocal repertoires and produce different types of vocalizations during different behavioral contexts. We searched for the most common adult vocalizations available in the repositories, and identified them as different types based on the visual inspection of the spectrograms and on the descriptions available from the Birds of the World database (59). For each species we analyzed one or two types of vocalizations. In songbirds (order Passeriformes) vocalizations are commonly categorized as either songs, which typically serve a sexual or territorial function, or calls, which are generally non-sexual and include contact, alarm and begging calls (60). While this distinction is clearly defined in most songbird species, it becomes ambiguous when applied to the vocalizations of other avian orders. Our dataset included a broad range of vocalization types (as described in the Xeno-Canto and Birds of the World databases) such as flight calls, alarm calls, contact calls, songs, and echolocation clicks (see the complete list of vocalizations we analyzed and their Xeno-Canto identifier in Dataset S1). Rather than attempting to impose the song versus call dichotomy or classifying vocalizations into narrow categories, we analyzed all vocalization types together in a single, integrated framework. Accordingly, we avoid referring to songs or calls throughout the article and instead use the general term “vocalizations”.

For each vocalization type we downloaded between 2 to 8 audio-files, and selected between 1 to 5 vocalizations from each file. We excluded from our search vocalizations produced by juveniles (begging calls) and sounds produced exclusively through non-vocal mechanisms (e.g., feather flutters, beak snapping or drumming). We included the bubbling display of the ruddy duck (*Oxyura jamaicensis*) into our dataset, as this display includes both a vocalization and sound produced by tapping the beak on the chest. We excluded two species with available audiogram data from our vocalizations data set. These were the common starling (*Sturnus vulgaris*), because of the difficulty to reliably analyze their extremely rich vocal repertoire that includes imitations of other birds, and the surf scoter (*Melanitta perspicillata*) which is considered generally silent (59), and for which we couldn’t find enough good-quality recordings of their vocalizations.

In total, we collected auditory and vocal data for 72 species of birds belonging to 15 different orders, and analyzed 108 different types of vocalizations for these species. These correspond to 0.64% of the total number of bird species (72 out of 11,276 species) and 34% of total number of bird orders (15 out of 44 orders) recognized by the International Ornithological Congress World Bird List (version 14.2, accessed on the 13^th^ of November 2024). The species included in our study represent the majority of those for which audiograms have been recorded through behavioral or physiological methods in adult individuals (excluding single-unit recordings or recordings at central level). We do not rule out the possibility that some species may have been missed in our search for audiogram data. We are, however, confident these will represent a minor fraction of species, and that their inclusion is unlikely to affect the main conclusions of this study.

The audio-files were transformed from Mp3 to Wav whenever necessary, and re-sampled to 22.05 kHz, with the exception of the vocalizations of the brown-headed cowbird (*Molothrus ater*), which can have frequency components that reach around 12kHz, and therefore these were resampled to 25 kHz. We manually selected the vocalizations from the oscillogram using RavenPro (61), taking care to only select units that were free from other unwanted noises, and that were clearly identifiable in the oscillogram and the spectrogram. The acoustic analysis of these manual selections was done in R using the libraries “seewave” (62) and “warbleR”(63). The audio files were high-pass filtered above 100 Hz to reduce the presence of low-frequency noises. For some low-frequency vocalizations (e.g., the hoots of many owls) we pass-filtered above 10 Hz to avoid modifying the structure of the vocalization. Additional filters were applied if deemed necessary during the visual inspection of the audio-files, taking care not to affect the frequency content of the vocalizations. We computed the power spectrum of each vocalization (windows size: 1024 samples, overlap: 90%, windowing function: Hanning), and averaged the spectra of each vocalization type and species to a single average spectrum. Average spectra were normalized to a maximum value of 0 dB. Averages and standard deviations of dB values were obtained using the *meandB()* and *sddB()* functions in the “seewave” library in R (62).

### Measuring auditory and vocal variables

We obtained three variables from each average power spectrum: a) the peak frequency (kHz), b) the high-frequency vocal limit (kHz), c) the low-frequency vocal limit (kHz). The peak frequency corresponds to the frequency with the highest energy of the power spectrum, and the high- and low-frequency limits were computed as the frequency of the spectrum 12 dB below the peak frequency (**Fig. 5A**). Our choice of using a 12 dB threshold was based on our evaluation of the quality of the audio files we analysed. Most of the recordings were made in the field with the animals typically located a few meters away from the recorder (in some cases, individuals were not even directly observed). Therefore, although we only analysed high-quality recordings, a higher threshold (e.g., at 20 or 30 dB) would have reduced our sample size and yielded less reliable bandwidth estimates (e.g., by falling into the background noise). Furthermore, we were interested in the fine tuning between vocalizations and hearing, and using a higher cut-off would have resulted in wider bandwidths that do not necessarily capture the bandwidth of the most prominent spectral components of the vocalizations.

**Figure 5.**
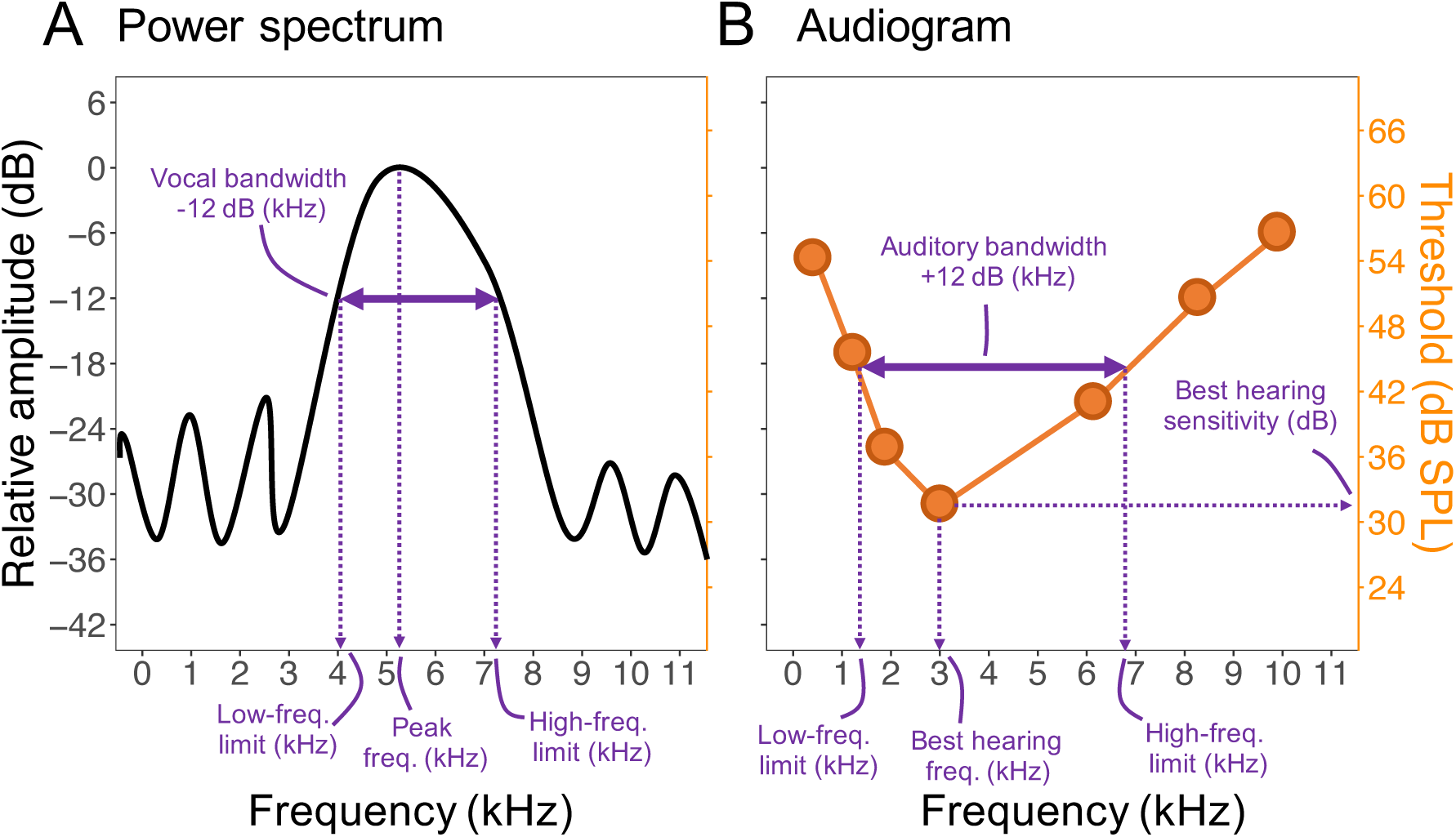
Diagrams showing a hypothetical A) power spectrum (black line) and B) audiogram (orange dots and line) together with the variables measured from each.

From each audiogram we obtained four variables: a) the best hearing frequency (kHz), b) the best hearing sensitivity (dB), c) the high-frequency hearing limit (kHz), and d) the low-frequency hearing limit (kHz). The best hearing frequency corresponds to the frequency value with the lowest amplitude threshold of the audiogram (i.e., the frequency value at the lowest point of the auditory curve), and the best hearing sensitivity corresponds to the threshold value at the best hearing frequency. The high- and low-frequency limits were computed as the threshold values at 12 dB above the best hearing frequency (**Fig. 5B**). Although a 30 dB cut-off is regularly used to define the auditory limits of birds(28) , we chose a 12 dB cut-off so we could have equivalent bandwidth measures for both the vocalizations and the audiogram (see explanation above).

### Comparing vocal and auditory data

The vocal and auditory variables were combined to compute three metrics that quantify their matching. First, we computed the difference between the peak frequency of the vocalizations and the best hearing frequency of the audiogram (**Fig. 1C**):

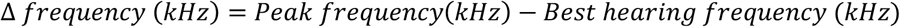

This measure reflects how far apart is the most prominent spectral component of the vocalization from the most sensitive point of the audiogram in terms of frequency. Positive values are indicative of species vocalizing at higher peak frequencies than their best hearing frequency, while negative values indicate species vocalizing with lower peak frequencies. A Δ frequency of exactly 0 kHz corresponds to a perfect match between the peak frequency and the best hearing frequency.

Second, we computed the difference between the auditory threshold estimated at the peak frequency of the vocalization and the best hearing threshold of the audiogram (**Fig. 1C**):

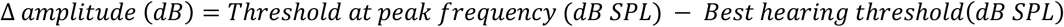

The threshold values at the peak frequency of the vocalizations were estimated through linear interpolation. Because this amplitude metric is computed relative to the best hearing threshold, its values are comparable between audiograms obtained through behavioral and electrophysiological methods. This measure can take only positive values and reflects how far from the best hearing frequency is the peak frequency of the vocalizations in terms of amplitude. Smaller values of Δ amplitude indicate better auditory sensitivity to the peak frequency of the vocalization, and a value of exactly 0 dB indicates a perfect match between peak frequency and best hearing frequency (i.e., a Δ amplitude value of exactly zero is unavoidably associated with a Δ amplitude value of exactly 0). The peak frequency of their vocalizations of a few species (*Phalacrocorax carbo*, *Somateria mollissima*, *Strix seloputo*, *Strix woodfordii*, *Ketupa zeylonensis*, *Bubo nipalensis*, *Bubo bubo* and *Asio otus*) fell below the minimum frequency tested in the audiogram, and therefore we could not interpolate the threshold at the peak frequency. In these cases, we used linear extrapolation to estimate the threshold value based on the two lowest frequencies for which auditory thresholds were available (***SI appendix***, **Figs. S5, S9, S17**).

Third, we calculated the overlap between the hearing and the vocal bandwidths (**Fig. 1C**). The bandwidths were defined as the range between the low- and high-frequency limits of vocalizations and audiograms. Some vocalizations presented secondary peaks with acoustic energy between 0 and -12 dB from the peak (e.g., vocalizations with harmonics). In these cases, we also included the bandwidths of the second and third most prominent peaks (the first peak is the peak frequency) in the calculation of the bandwidth overlaps. Bandwidth overlaps were computed in Hertz and transformed to percent relative to the hearing bandwidth using the following formula:

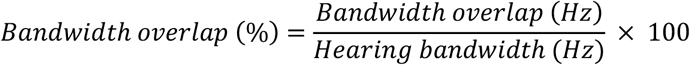

While Δ frequency and Δ amplitude are point estimates of vocal and auditory matching, the bandwidth overlap quantifies the correspondence between the most prominent vocal elements and the region of best audition in terms of ranges. A 0% bandwidth overlap indicates a complete mismatch between the vocal and the hearing bandwidths, and 100% bandwidth overlap indicates that the vocalization bandwidth fully occupies the hearing bandwidth.

### Body weight data

For all the 72 species we collected body weight (gr) data from the AVONET database (64). In the database, body mass data corresponds to the average across males and females, therefore we did not consider sexual dimorphism in our analyses, although we assume that intra-specific variation in weight is smaller than the inter-specific variation we were interested in. Body weight data was log10-transformed for all the analyses.

### Phylogenetic tree

We downloaded 1,000 phylogenetic trees (with Hackett backbone) containing the 72 species in the dataset from the BirdTree website (65). These trees are sampled from a pseudo-posterior distribution after trimming a subset of taxa from a larger phylogeny of birds. The northern white-faced owl (*Ptilopsis leucotis*), the black scoter (*Melanitta americana*) and the Carolina chickadee (*Poecile carolinensis*) were found in the trees under the names *Otus leucotis*, *Melanitta nigra* and *Parus carolinensis*, respectively. The 1,000 trees were summarized into a single maximum clade credibility (MCC) tree with median node heights using TreeAnnotator (66), and this summary tree was used for performing phylogenetic comparative analyses (**Fig. 1A**).

### Phylogenetic analyses

All the comparative phylogenetic analyses were run in R using the library “brms” (67) to fit Bayesian generalized phylogenetic linear models. The figure of the MCC phylogeny was created using the library “phytools” (68). We computed the variance-covariance matrix of the MCC phylogeny assuming a Brownian motion model of evolution using the library “ape” (69), and included this matrix as a random factor in all the models to account for phylogenetic relatedness. For half of the species in the data set (N = 36) we analysed two different types of vocalizations, and therefore we included a random intercept for each species to account for these repeated measures. Models that excluded repeated measures excluded this random effect. For all the models we ran 6 chains of 10,000 iterations each with a warm up of 4,000 iterations, and used the default weakly informative priors from “brms”. These priors improve model convergence and have small influence on parameter estimation (67).

### Phylogenetic signal

We estimated Pagel’s λ for all the vocal and auditory traits, as well as body mass using the “phytools” package (68). For species with two types of vocalizations, we calculated the mean value and assessed the phylogenetic signal based on this average.

### Analysis of auditory and vocal variables

First, we tested whether the auditory variables (best frequency, low- and high-frequency hearing limits) and vocal variables (peak frequency, low- and high-frequency vocal limits) were allometrically related to body weight. For each variable we fitted a separate model and included body weight as their only predictor. For the auditory variables, we chose an inverse Gaussian distribution and logarithmic link function. For the vocal variables, we chose a Gaussian distribution and logarithmic link function.

Then we evaluated if auditory and vocal variables were related to each other. For this we fitted three different models. A first model included peak frequency as the response and best hearing frequency as predictor, a second model included low-frequency vocal limit as the response and low-frequency hearing limit as predictor, and a third model included high-frequency vocal limit as the response and high-frequency hearing limit as predictor. In these three models we added body weight as a covariate, and assumed a Gaussian distribution for the response and a logarithmic link function.

### Analysis of vocal and auditory matching

First, we quantified general trends of vocal and auditory matching across all birds. For this, we fitted three separate phylogenetic mixed models (species as random effect, see above), including Δ frequency (kHz), Δ amplitude (dB) and bandwidth overlap (%) as response variable, and only the intercept as predictor (i.e., a model with formula: y ∼ 1). For the analysis of Δ frequency we chose a Gaussian distribution with identity link function, for Δ amplitude we chose a Gamma distribution with logarithmic link function, and for bandwidth overlap we transformed the data from percent to proportions (between 0 and 1) and chose a zero-one inflation Beta distribution with logit link function for the mean parameter.

Next, we evaluated if our three measures of auditory-vocal matching (i.e., Δ frequency, Δ amplitude and bandwidth overlap) allometrically scaled with body weight. For this we included body weight as the only predictor in the model. For the analysis of Δ frequency we used a Gaussian distribution with identity link function, for Δ amplitude we used a Gamma distribution with a logarithmic link function and for bandwidth overlap (as proportion) we used a zero-one inflation beta distribution with logit link function for the mean parameter.

For all the models we assessed chain convergence and stability by ensuring that 𝑅^J^ (r-hat) values were not above 1.01 and by visually inspecting trace plots. Model fit was visually evaluated using posterior predictive graphs. We report the mean of the posterior distributions of the coefficients estimated by the models accompanied by their corresponding credible intervals (CI). We report the 95% CI in most cases, and in a few occasions report the 89% CI to indicate weak evidence for an effect. Both 95 and 89% CIs are arbitrary thresholds (70), but the 89% CI is considered to be more stable (71). We evaluated the evidence for the presence of an effect and its direction based on the mean of the posterior distribution and the credible intervals. Bayesian analyses aim to move away from relying on frequentist *P*-values to assess the existence of effects (72). However, *P*-values may still be helpful for readers unfamiliar with Bayesian statistics to interpret our results. We therefore computed the probability of direction for our model estimates (i.e., the probability of the effect is positive or negative) and transformed them to *P*-values using the library “bayestestR” in R (73).

## Acknowledgments

We are thankful to Jacintha Ellers for providing valuable comments on a previous version of the manuscript, and to all those people who have deposited their bird audio recordings to public repositories. MM was supported by BecasChile 2018-Comisión Nacional de Investigación y Desarrollo (CONICYT) scholarship (number: 72190501).

