## Supplementary material for "Optimal tuning not required: birds communicate effectively with mismatched auditory filters": SI Appendix

**This PDF file includes:**

Figures S1 to S18
Table S1
SI References

**Other supporting materials for this manuscript include the following:**

Dataset S1

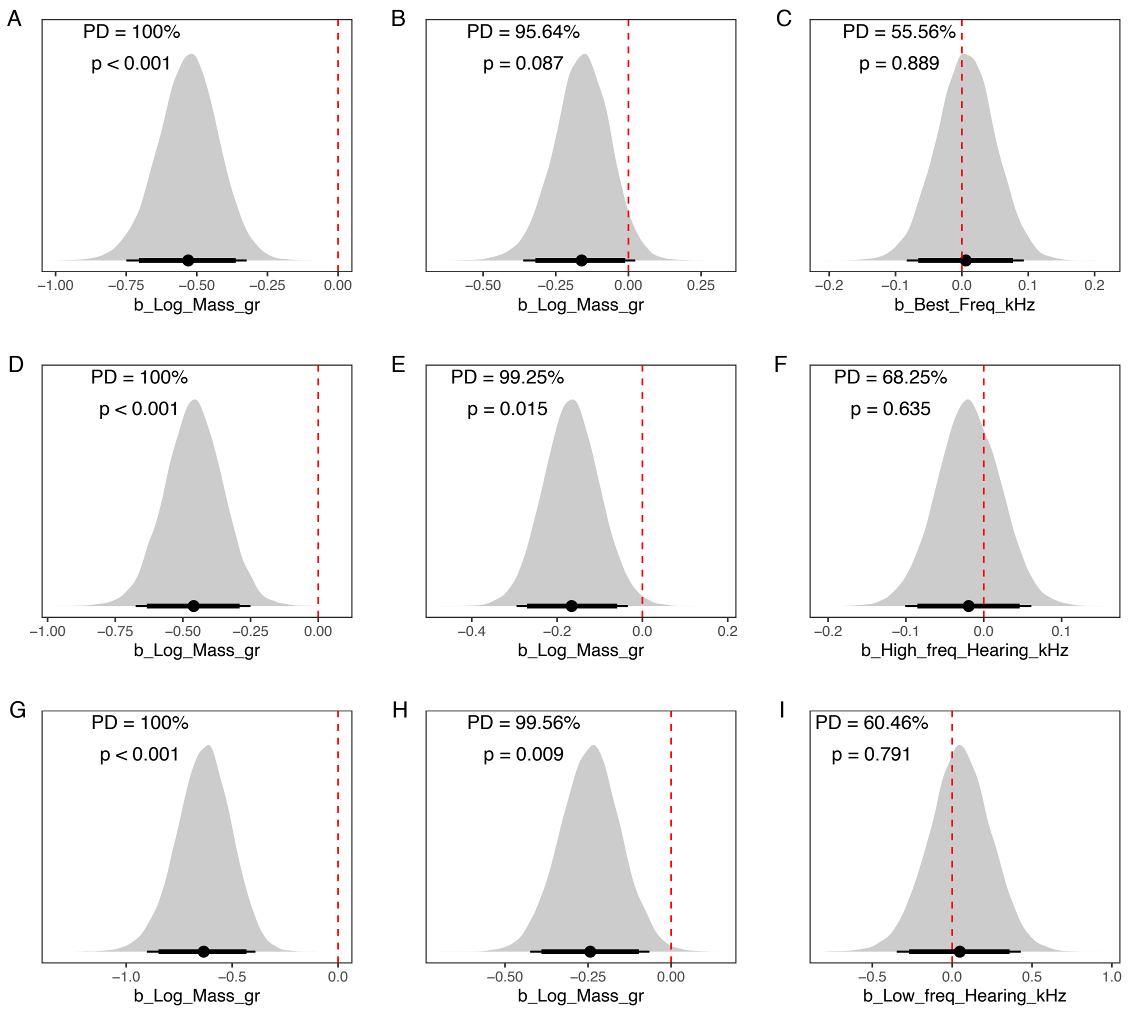


**Fig. S1.** Posterior distributions for the slope estimates of the linear models shown in Fig. 2 of the main text. The black point corresponds to the mean of the posterior distribution, and the horizontal bars show the 89 and 95% credible intervals. The probability of direction (PD) and the p-value of each posterior distribution are shown inside each panel.

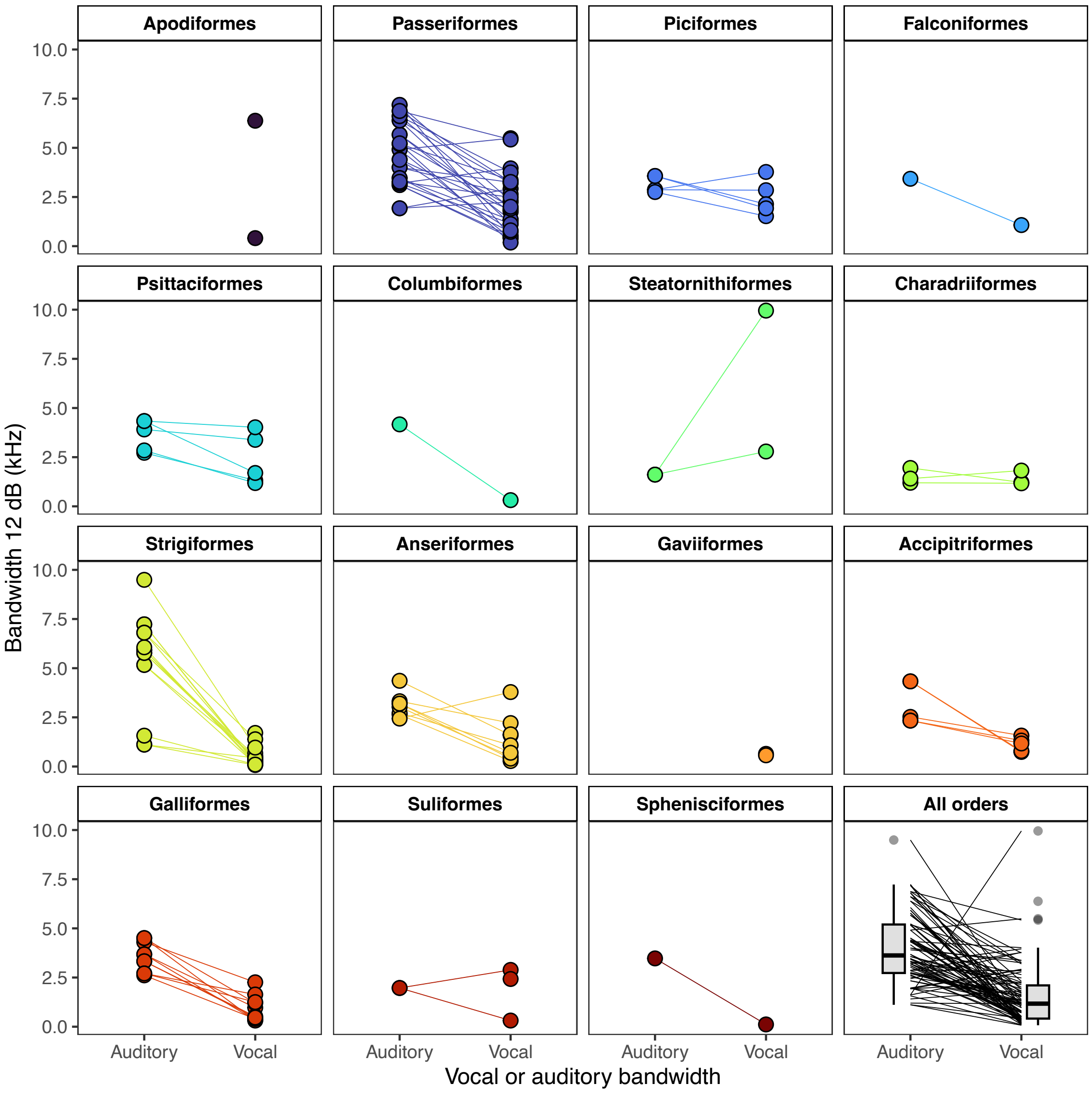


**Fig. S2.** Comparison between the auditory and vocal bandwidths. The different panels show different orders, and the horizontal lines are connecting species. When an auditory bandwidth connects to two vocal bandwidths is because two vocalizations were analysed for that species. For the vocalizations we used the bandwidth of the main peak of the spectrogram only. For Apodiformes and Gaviiformes bandwidths could not be computed because the audiograms did not reach 12 dB from the best hearing frequency in the low-frequency hearing limit. The panel at the bottom right shows all the species grouped together.

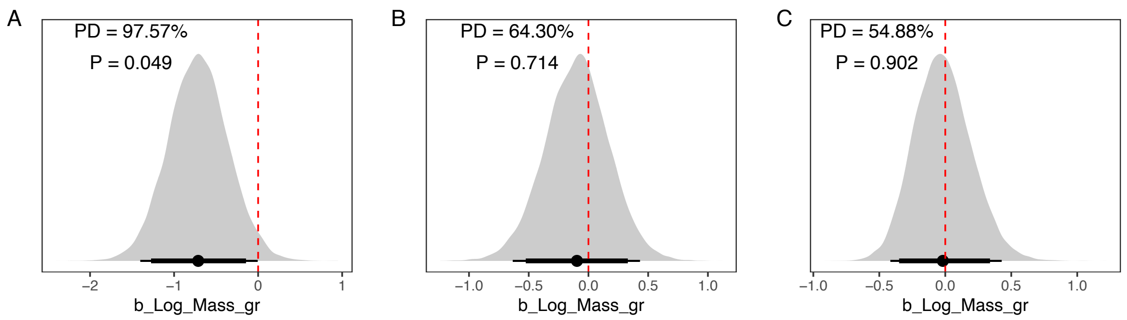


**Fig. S3**. Posterior distributions for the slope estimates of the linear models shown in Fig. 4 of the main text. The black point corresponds to the mean of the posterior distribution, and the horizontal bars show the 89 and 95% credible intervals. The probability of direction (PD) and the p-value of each posterior distribution are shown inside each panel**.**

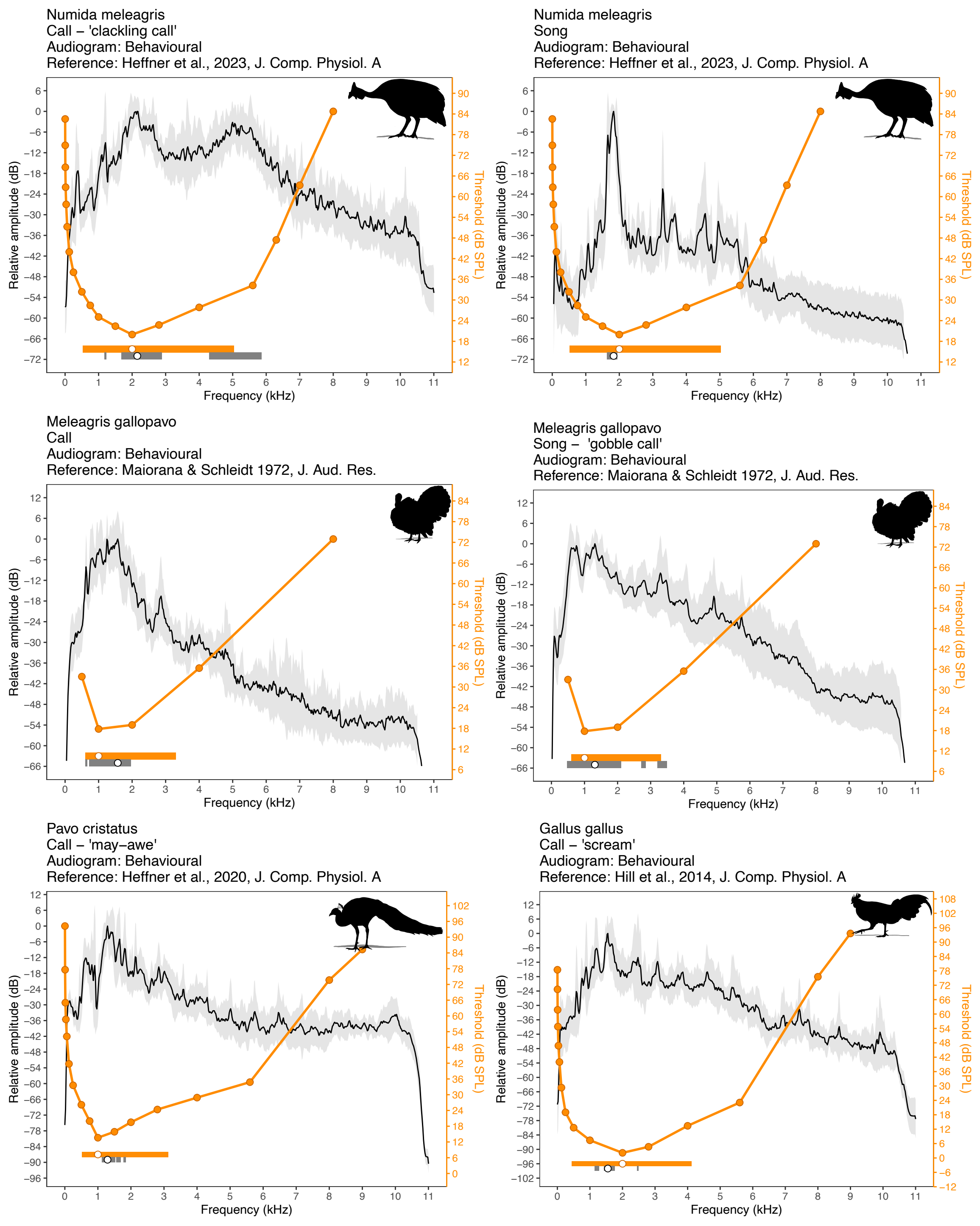


**Fig. S4:** Audiograms (orange) and mean power spectrum (black) for the order Galliformes. They grey shading corresponds to the S.D. of the power spectra analyzed. Horizontal bars at the bottom show the auditory (orange) and vocal (grey) bandwidths (at 12 dB), and the points inside of them depict the best hearing frequency of the audiogram and the peak frequency of the mean spectrum.

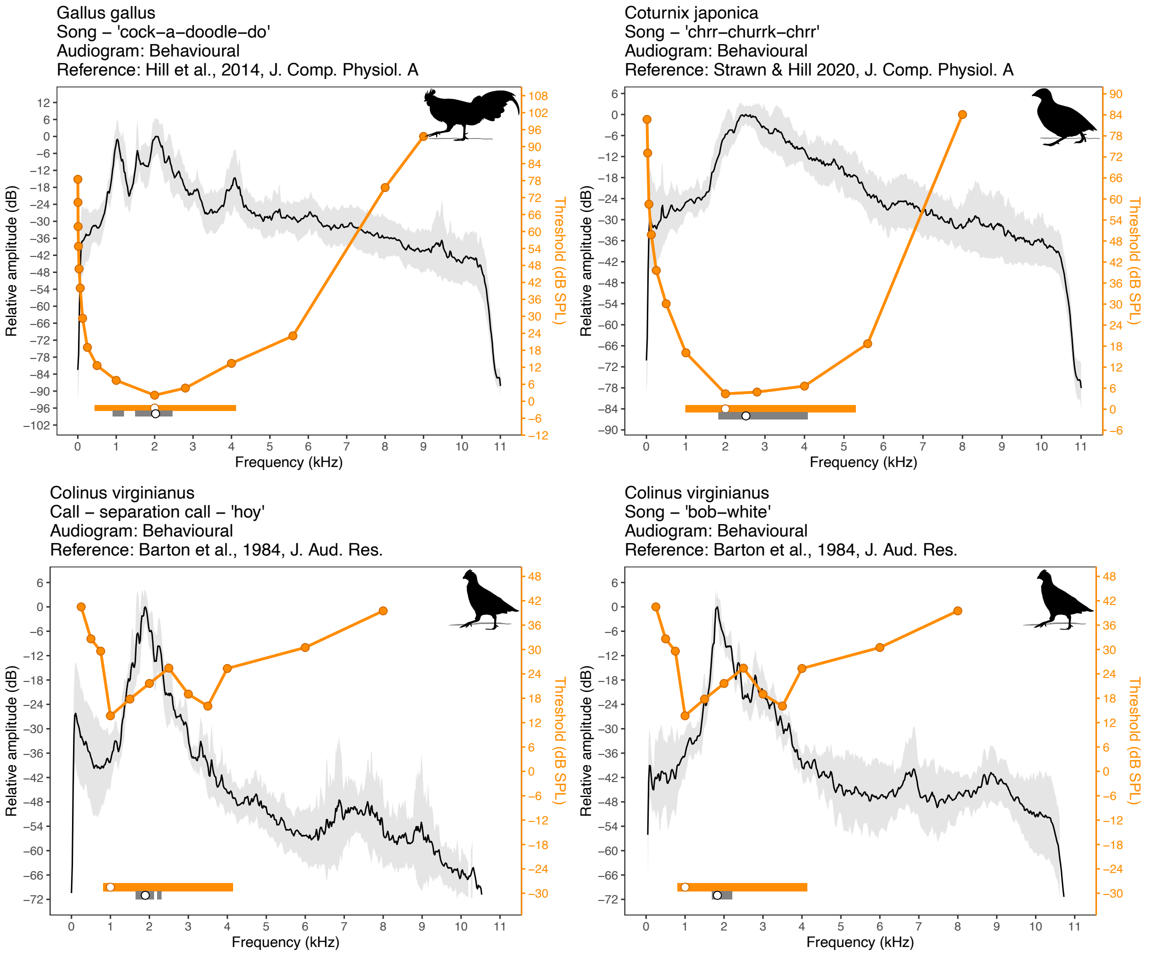


***(continued)*** **Fig. S4:** Audiograms (orange) and average power spectrum (black) for the order Galliformes. They grey shading corresponds to the S.D. of the power spectra analyzed. Horizontal bars at the bottom show the auditory (orange) and vocal (grey) bandwidths (at 12 dB), and the points inside of them depict the best hearing frequency of the audiogram and the peak frequency of the mean spectrum.

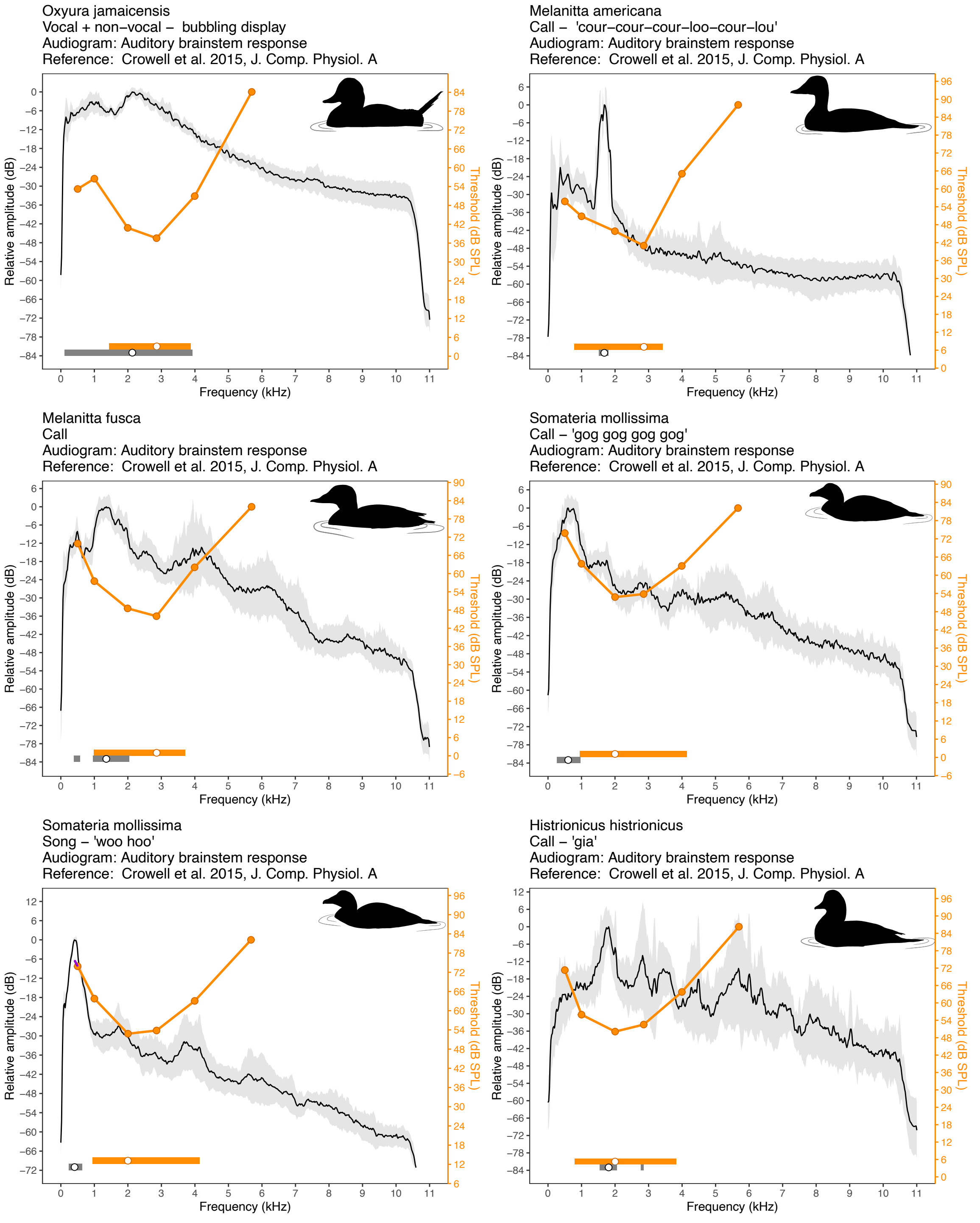


**Fig. S5:** Audiograms (orange) and average power spectrum (black) for the order Anseriformes. They grey shading corresponds to the S.D. of the power spectra analyzed. Horizontal bars at the bottom show the auditory (orange) and vocal (grey) bandwidths (at 12 dB), and the points inside of them depict the best hearing frequency of the audiogram and the peak frequency of the mean spectrum. The purple line in the audiogram of *Somateria mollissima* shows the linear extrapolation used to estimate the threshold at the peak frequency of its song.

Clangula hyemalis
Call – 'ahr–ahr–ahroulit'
Audiogram: Auditory brainstem response
Reference: Crowell et al. 2015, J. Comp. Physiol. A

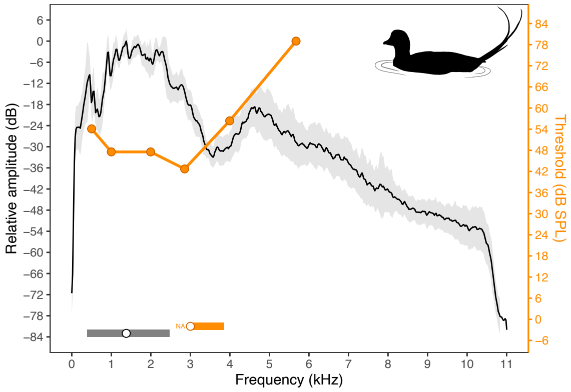


Aythya affinis
Call
Audiogram: Behavioural
Reference: Crowell et al., 2016, JASA

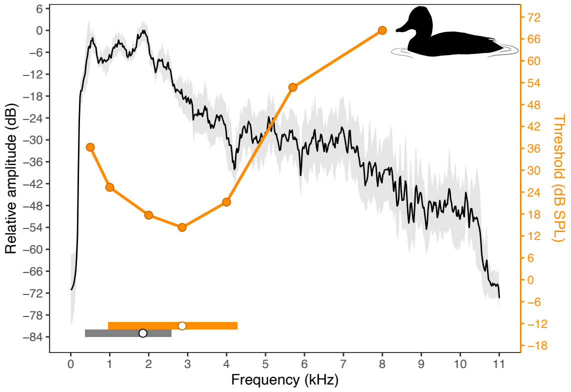


Anas platyrhynchos
CII – quacking call
Audiogram: Behavioural
Reference: Hill 2017, J. Comp. Physiol. A

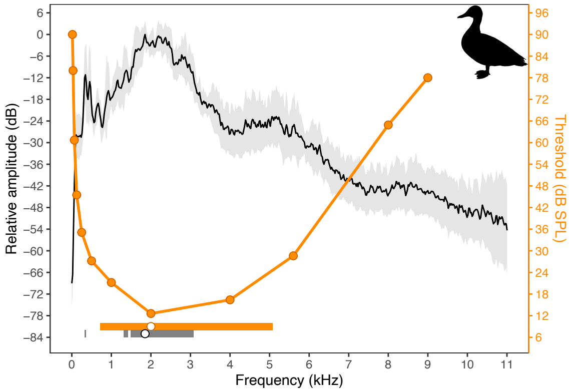


***(continued)*** **Fig. S5:** Audiograms (orange) and average power spectrum (black) for the order Anseriformes. They grey shading corresponds to the S.D. of the power spectra analyzed. Horizontal bars at the bottom show the auditory (orange) and vocal (grey) bandwidths (at 12 dB), and the points inside of them depict the best hearing frequency of the audiogram and the peak frequency of the mean spectrum.

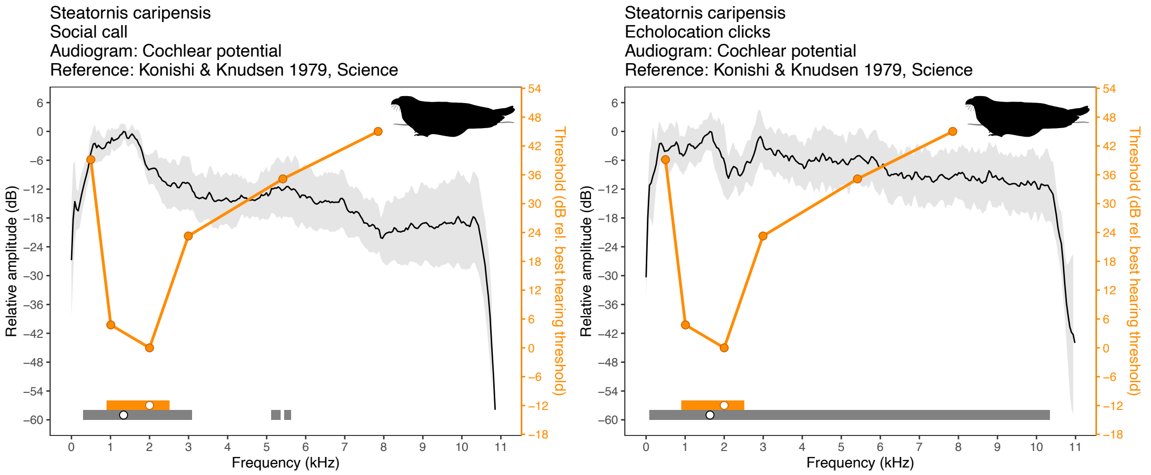


**Fig. S6:** Audiograms (orange) and average power spectrum (black) for the order Steatornithiformes. They grey shading corresponds to the S.D. of the power spectra analyzed. Horizontal bars at the bottom show the auditory (orange) and vocal (grey) bandwidths (at 12 dB), and the points inside of them depict the best hearing frequency of the audiogram and the peak frequency of the mean spectrum.

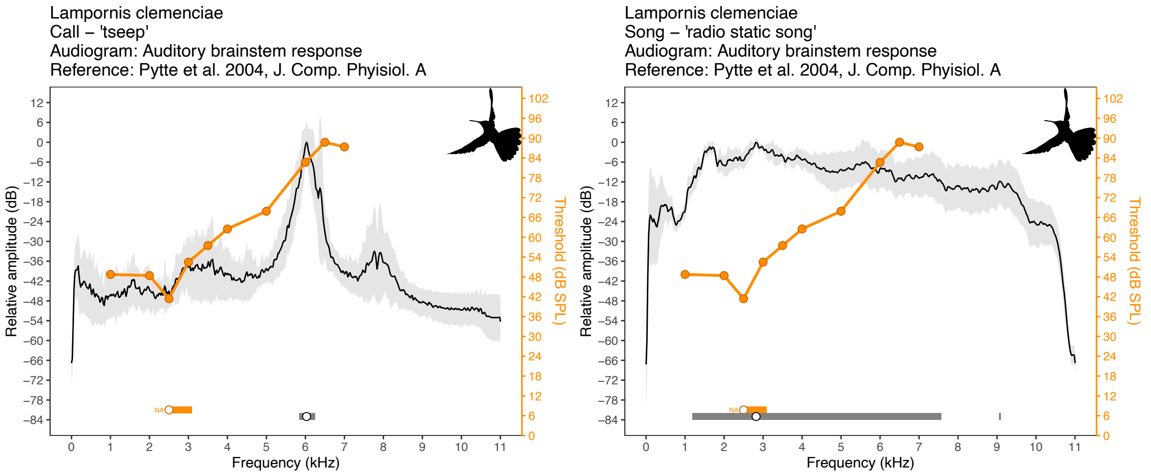


**Fig. S7:** Audiograms (orange) and average power spectrum (black) for the order Apodiformes. They grey shading corresponds to the S.D. of the power spectra analyzed. Horizontal bars at the bottom show the auditory (orange) and vocal (grey) bandwidths (at 12 dB), and the points inside of them depict the best hearing frequency of the audiogram and the peak frequency of the mean spectrum.

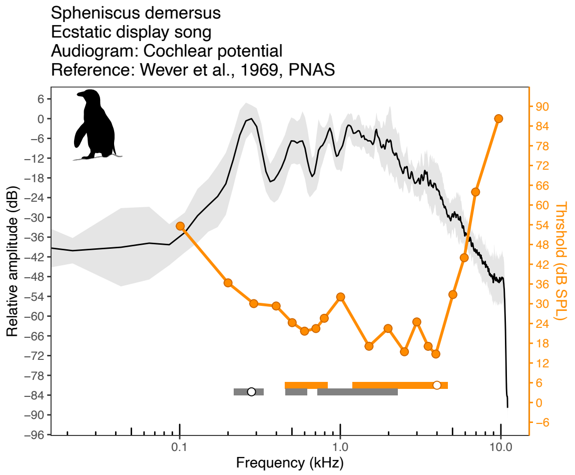


**Fig. S8:** Audiograms (orange) and average power spectrum (black) for the order Sphenisciformes. They grey shading corresponds to the S.D. of the power spectra analyzed. Horizontal bars at the bottom show the auditory (orange) and vocal (grey) bandwidths (at 12 dB), and the points inside of them depict the best hearing frequency of the audiogram and the peak frequency of the mean spectrum.

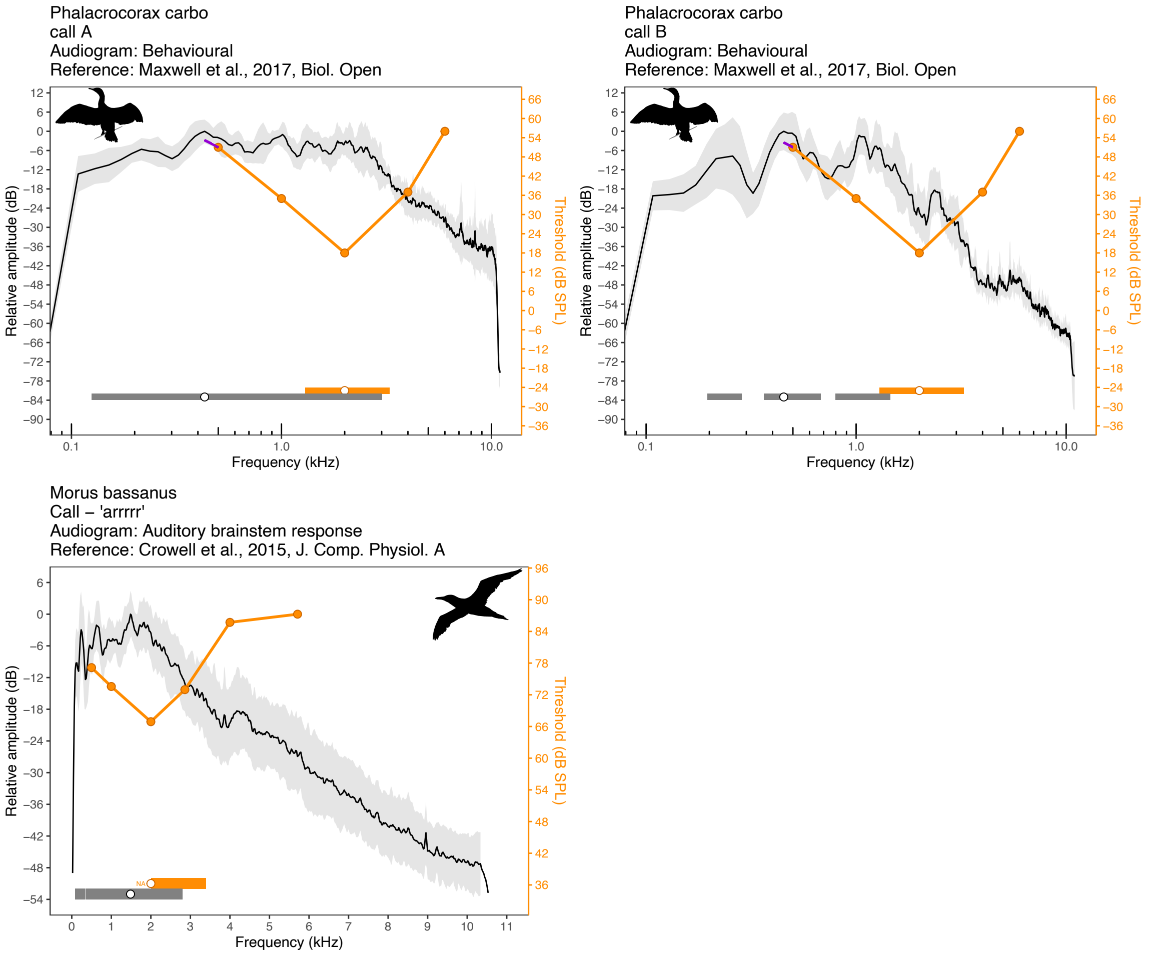


**Fig. S9:** Audiograms (orange) and average power spectrum (black) for the order Suliformes. They grey shading corresponds to the S.D. of the power spectra analyzed. Horizontal bars at the bottom show the auditory (orange) and vocal (grey) bandwidths (at 12 dB), and the points inside of them depict the best hearing frequency of the audiogram and the peak frequency of the mean spectrum. The purple lines in the audiograms of *Phalacrocorax carbo* show the linear extrapolation used to estimate the threshold at the peak frequency of both types of calls.

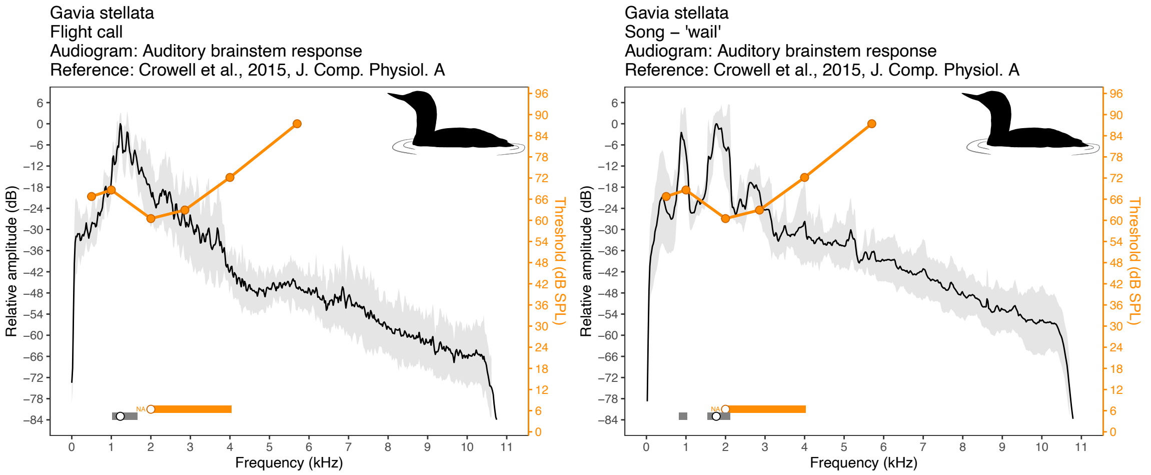


**Fig. S10:** Audiograms (orange) and average power spectrum (black) for the order Gaviiformes. They grey shading corresponds to the S.D. of the power spectra analyzed. Horizontal bars at the bottom show the auditory (orange) and vocal (grey) bandwidths (at 12 dB), and the points inside of them depict the best hearing frequency of the audiogram and the peak frequency of the mean spectrum.

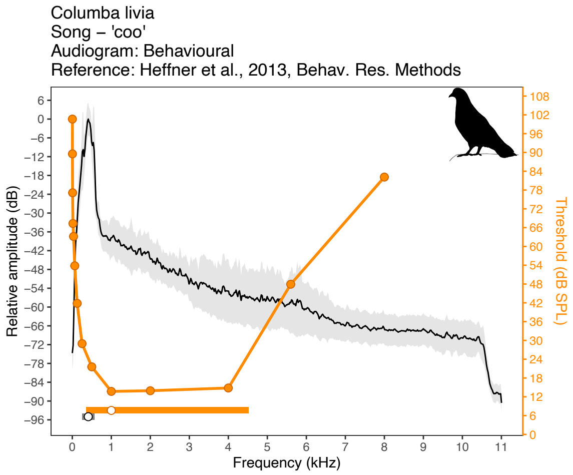


**Fig. S11:** Audiograms (orange) and average power spectrum (black) for the order Columbiformes. They grey shading corresponds to the S.D. of the power spectra analyzed. Horizontal bars at the bottom show the auditory (orange) and vocal (grey) bandwidths (at 12 dB), and the points inside of them depict the best hearing frequency of the audiogram and the peak frequency of the mean spectrum.

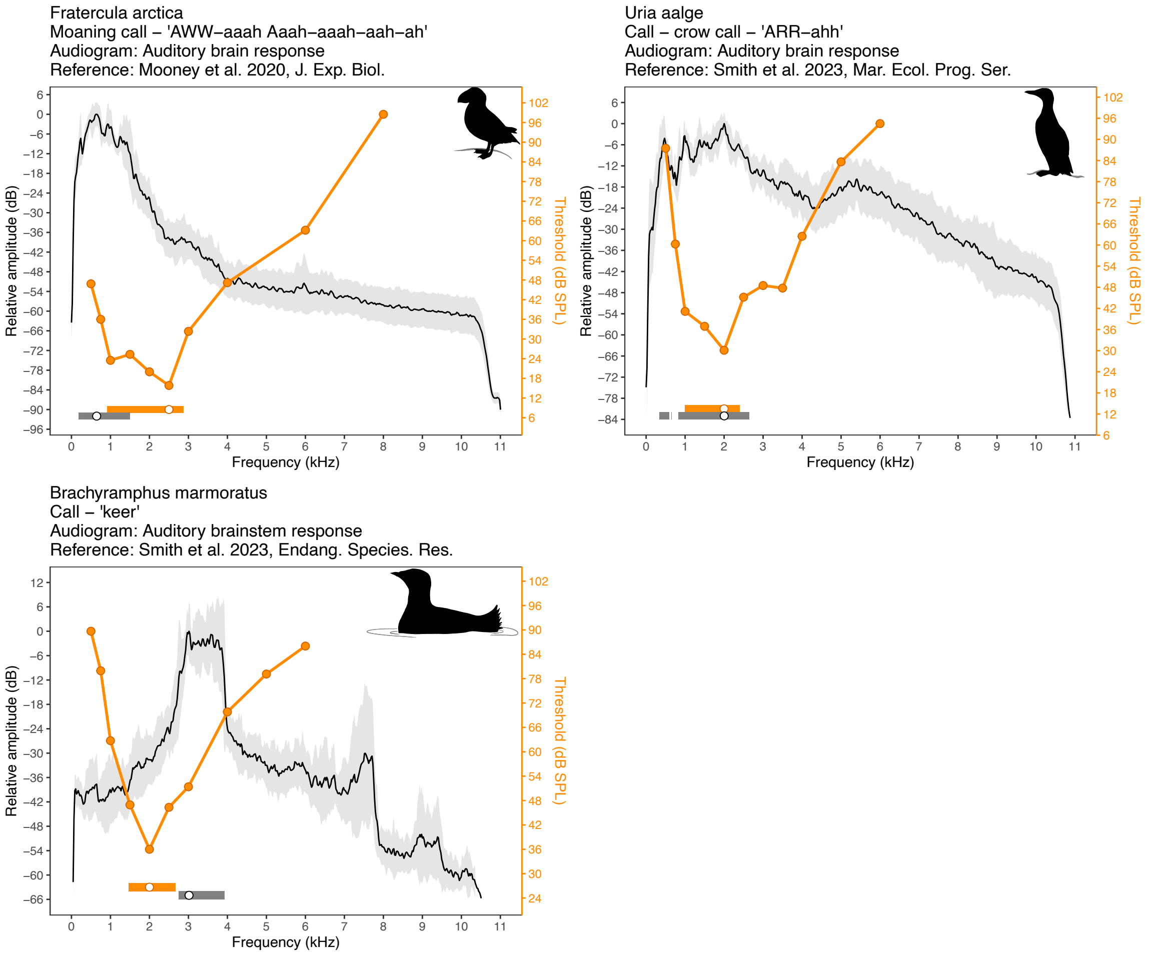


**Fig. S12:** Audiograms (orange) and average power spectrum (black) for the order Charadriiformes. They grey shading corresponds to the S.D. of the power spectra analyzed. Horizontal bars at the bottom show the auditory (orange) and vocal (grey) bandwidths (at 12 dB), and the points inside of them depict the best hearing frequency of the audiogram and the peak frequency of the mean spectrum.

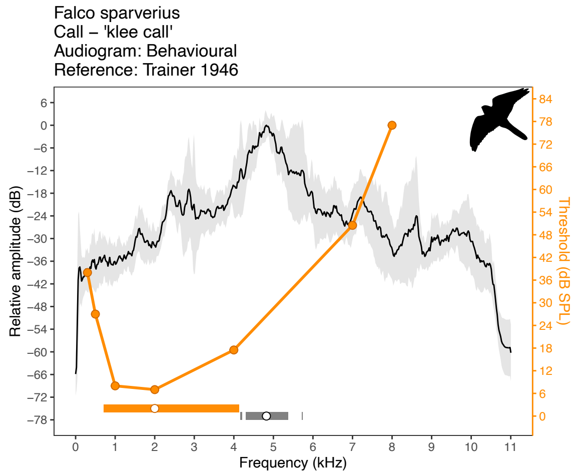


**Fig. S13:** Audiograms (orange) and average power spectrum (black) for the order Falconiformes. They grey shading corresponds to the S.D. of the power spectra analyzed. Horizontal bars at the bottom show the auditory (orange) and vocal (grey) bandwidths (at 12 dB), and the points inside of them depict the best hearing frequency of the audiogram and the peak frequency of the mean spectrum.

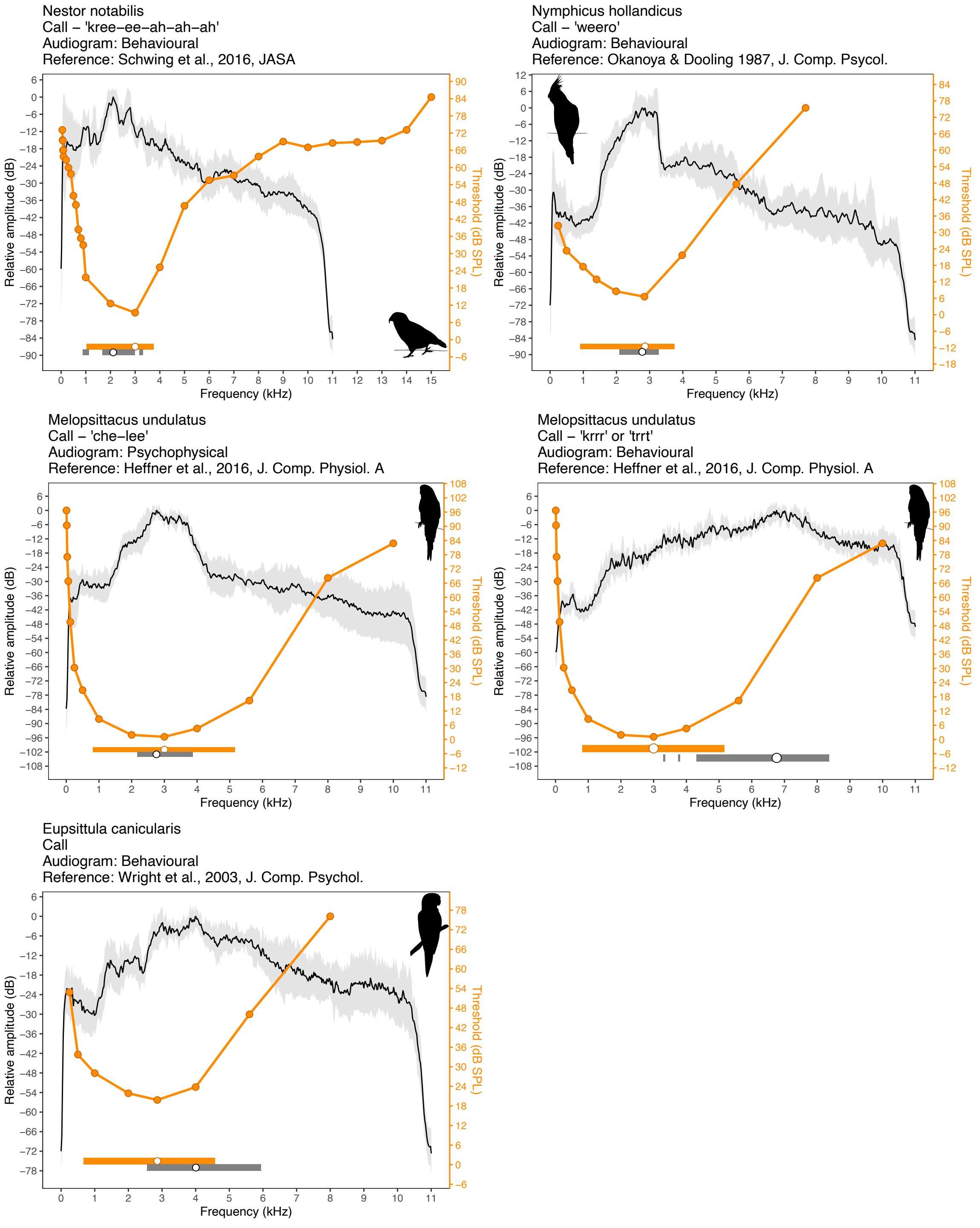


**Fig. S14:** Audiograms (orange) and average power spectrum (black) for the order Psittaciformes. They grey shading corresponds to the S.D. of the power spectra analyzed. Horizontal bars at the bottom show the auditory (orange) and vocal (grey) bandwidths (at 12 dB), and the points inside of them depict the best hearing frequency of the audiogram and the peak frequency of the mean spectrum.

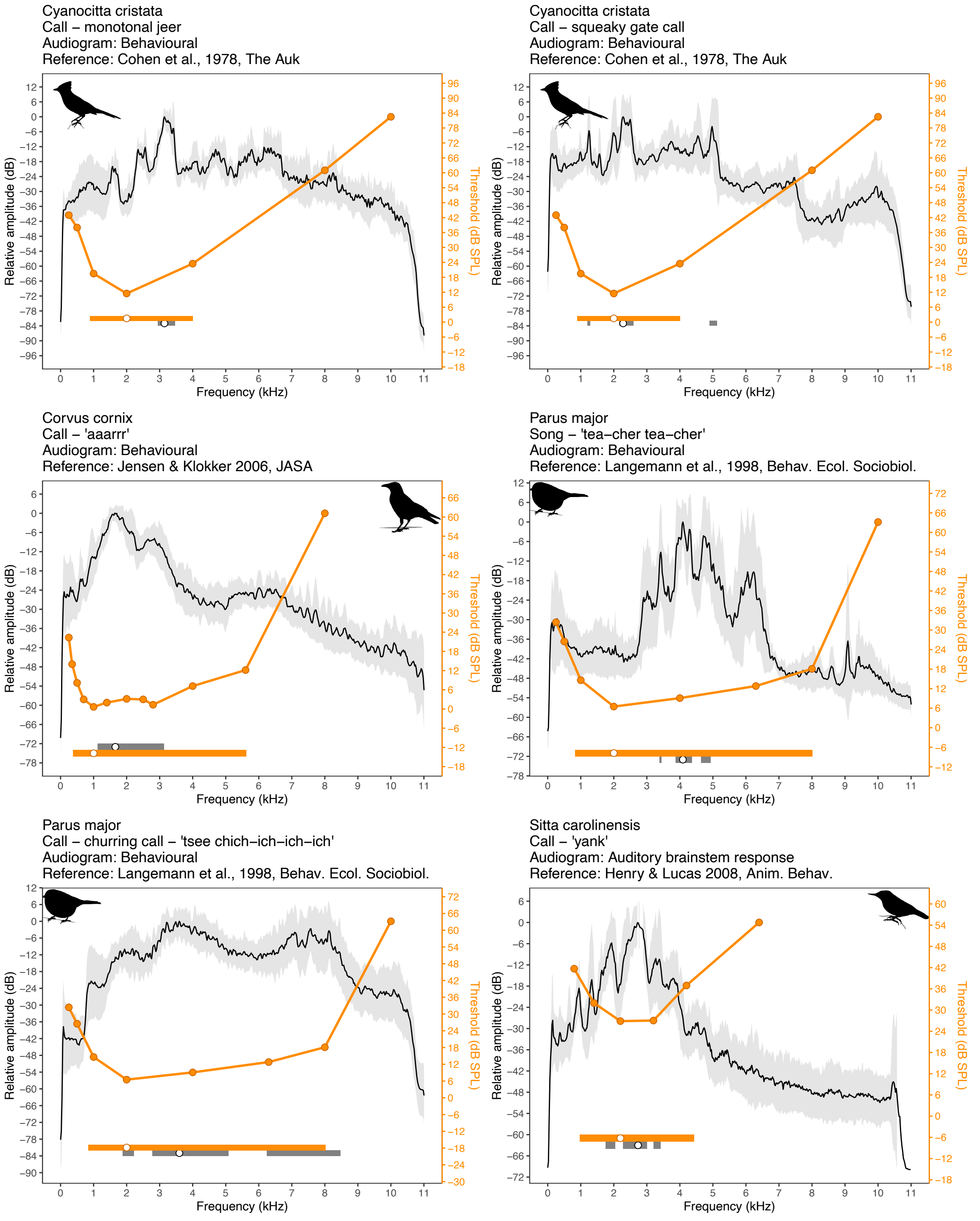


**Fig. S15:** Audiograms (orange) and average power spectrum (black) for the order Passeriformes. They grey shading corresponds to the S.D. of the power spectra analyzed. Horizontal bars at the bottom show the auditory (orange) and vocal (grey) bandwidths (at 12 dB), and the points inside of them depict the best hearing frequency of the audiogram and the peak frequency of the mean spectrum.

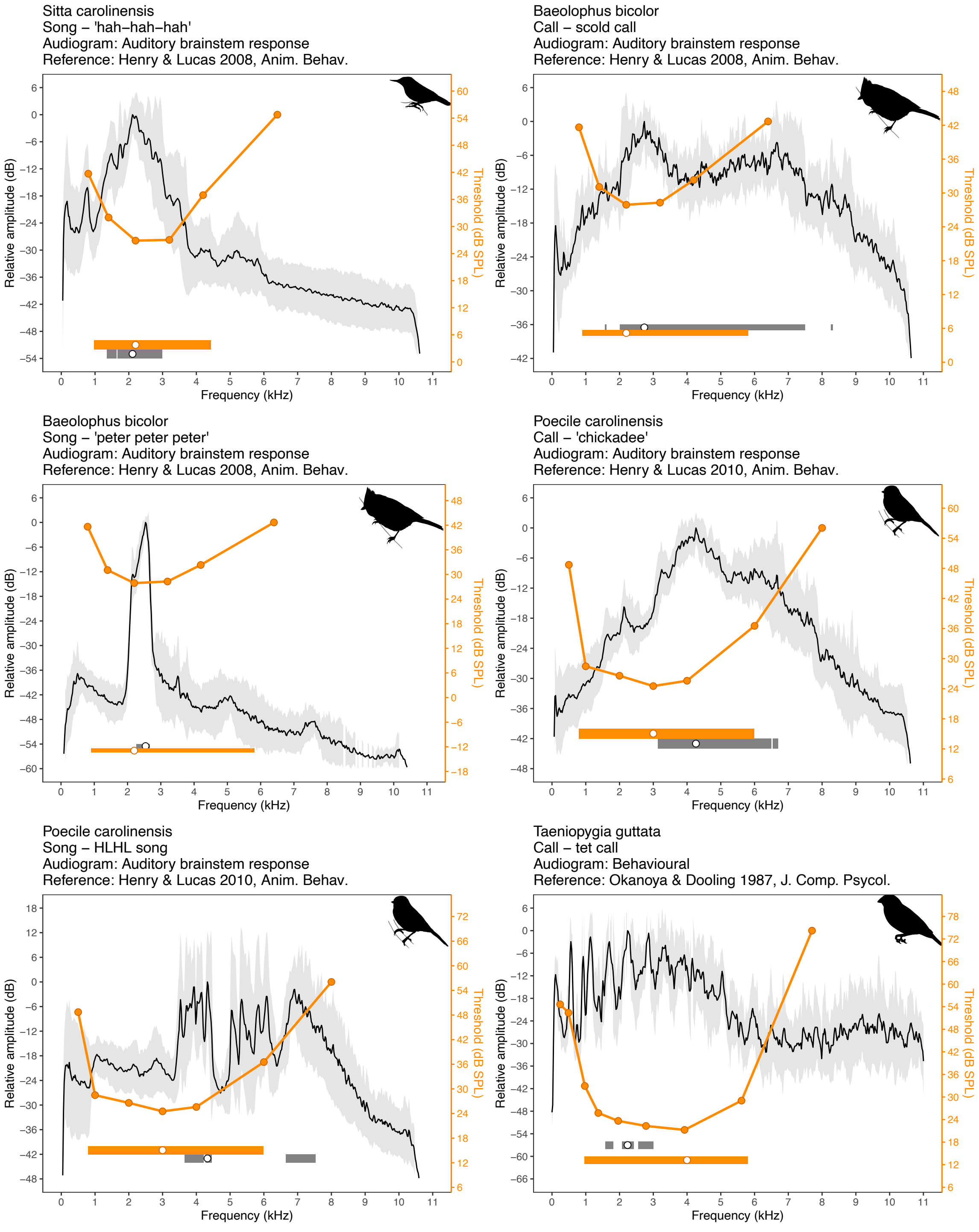


***(continued)*** **Fig. S15:** Audiograms (orange) and average power spectrum (black) for the order Passeriformes. They grey shading corresponds to the S.D. of the power spectra analyzed. Horizontal bars at the bottom show the auditory (orange) and vocal (grey) bandwidths (at 12 dB), and the points inside of them depict the best hearing frequency of the audiogram and the peak frequency of the mean spectrum.

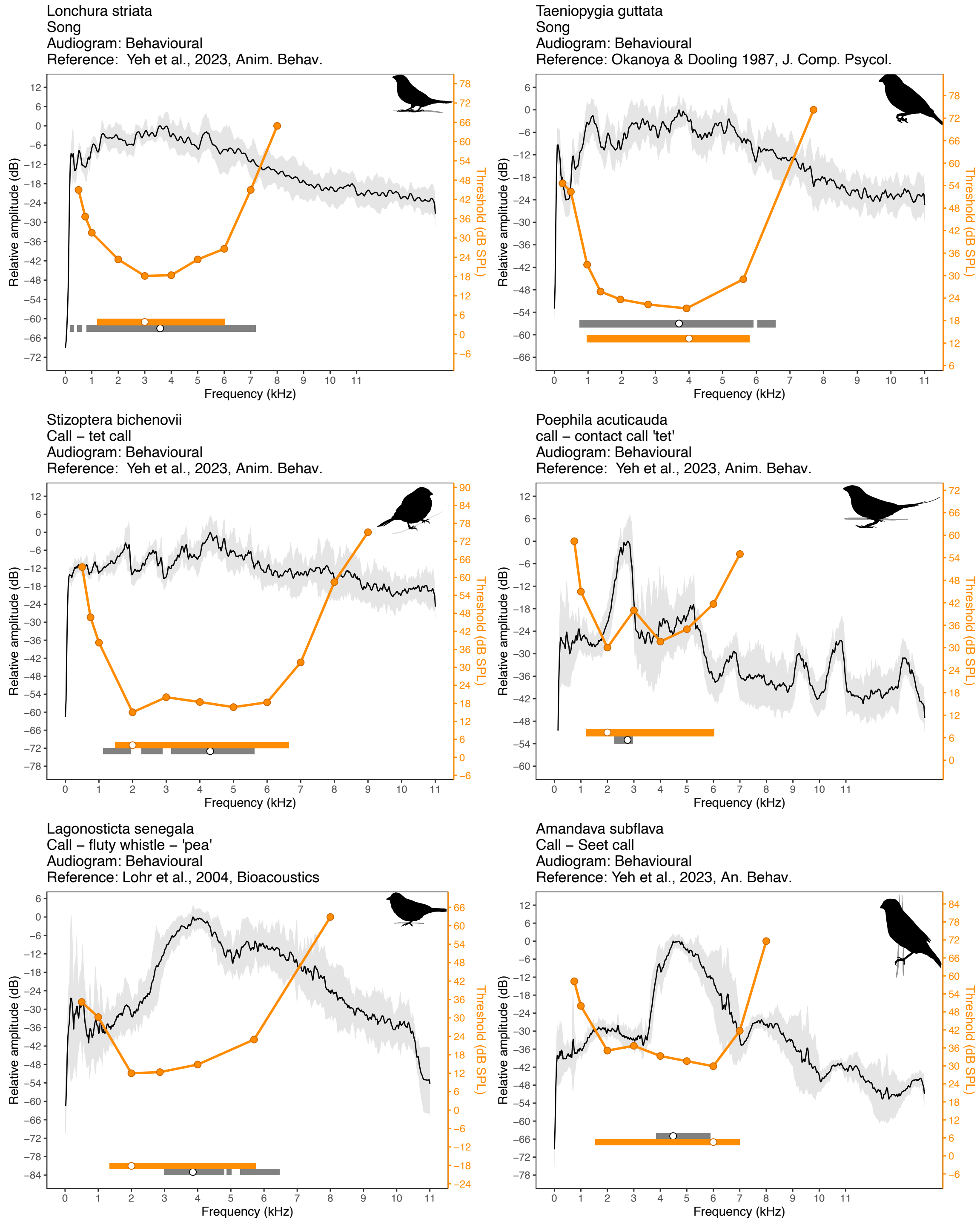


***(continued)*** **Fig. S15:** Audiograms (orange) and average power spectrum (black) for the order Passeriformes. They grey shading corresponds to the S.D. of the power spectra analyzed. Horizontal bars at the bottom show the auditory (orange) and vocal (grey) bandwidths (at 12 dB), and the points inside of them depict the best hearing frequency of the audiogram and the peak frequency of the mean spectrum.

Passer domesticus
Call– chatter?
Audiogram: Auditory brainstem response
Reference: Henry & Lucas 2008, Anim. Behav.

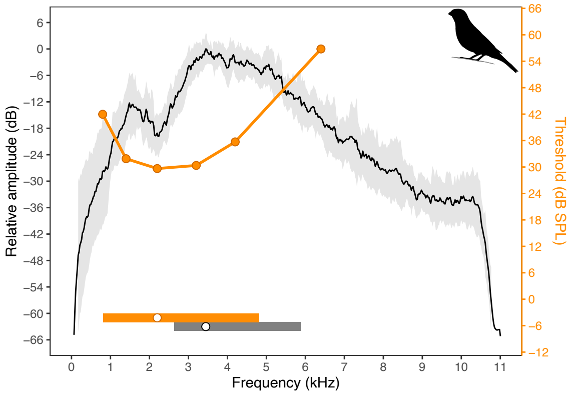


Passer domesticus
Song – 'cheep'
Audiogram: Auditory brainstem response
Reference: Henry & Lucas 2008, Anim. Behav.

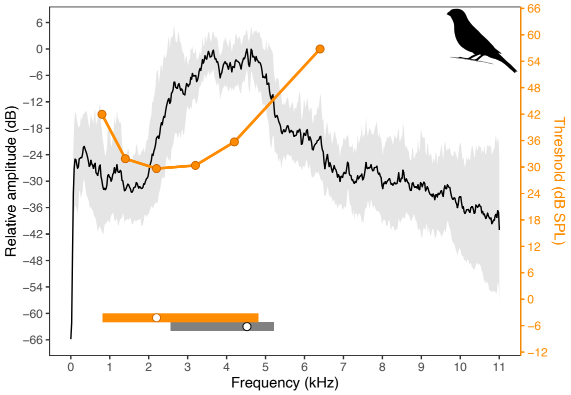


Serinus canaria
Song
Audiogram: Behavioural
Reference: Dooling et al., 1971, JASA

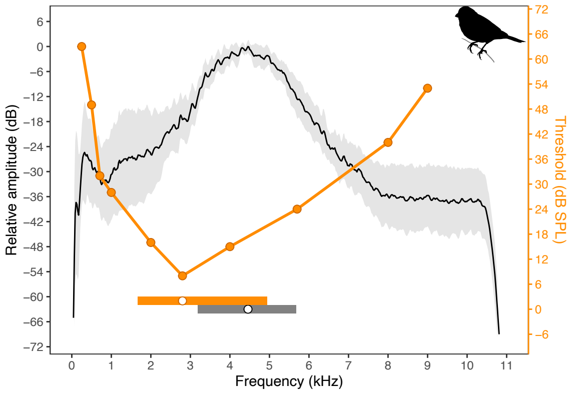


Pyrrhula pyrrhula
Call – 'peu'
Audiogram: Behavioural
Reference: Schwartzkopff 1949, Zeitschr. vergl. Physiol.

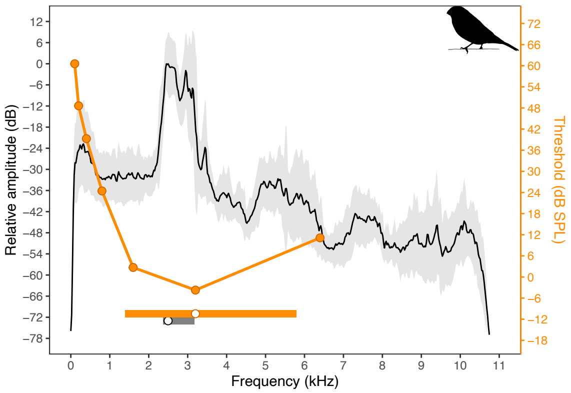


Haemorhous mexicanus
Call
Audiogram: Behavioural
Reference: Dooling et al. 1978, J. Comp. Physiol. Psycol.

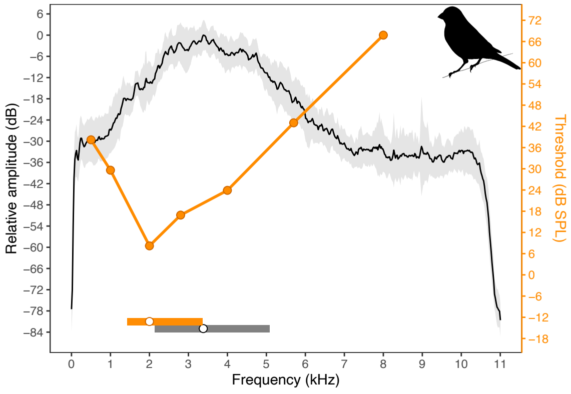


Haemorhous mexicanus
Song
Audiogram: Behavioural
Reference: Dooling et al. 1978, J. Comp. Physiol. Psycol.

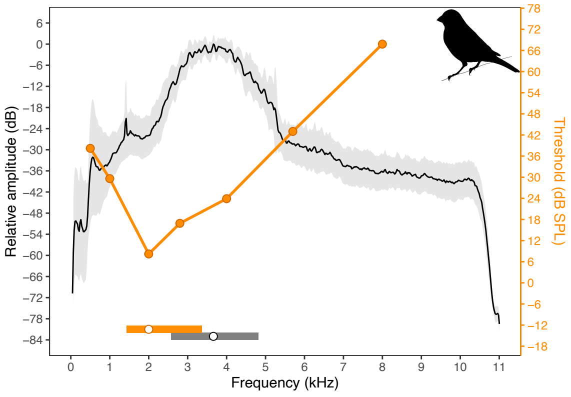


***(continued)* Fig. S15:** Audiograms (orange) and average power spectrum (black) for the order Passeriformes. They grey shading corresponds to the S.D. of the power spectra analyzed. Horizontal bars at the bottom show the auditory (orange) and vocal (grey) bandwidths (at 12 dB), and the points inside of them depict the best hearing frequency of the audiogram and the peak frequency of the mean spectrum.

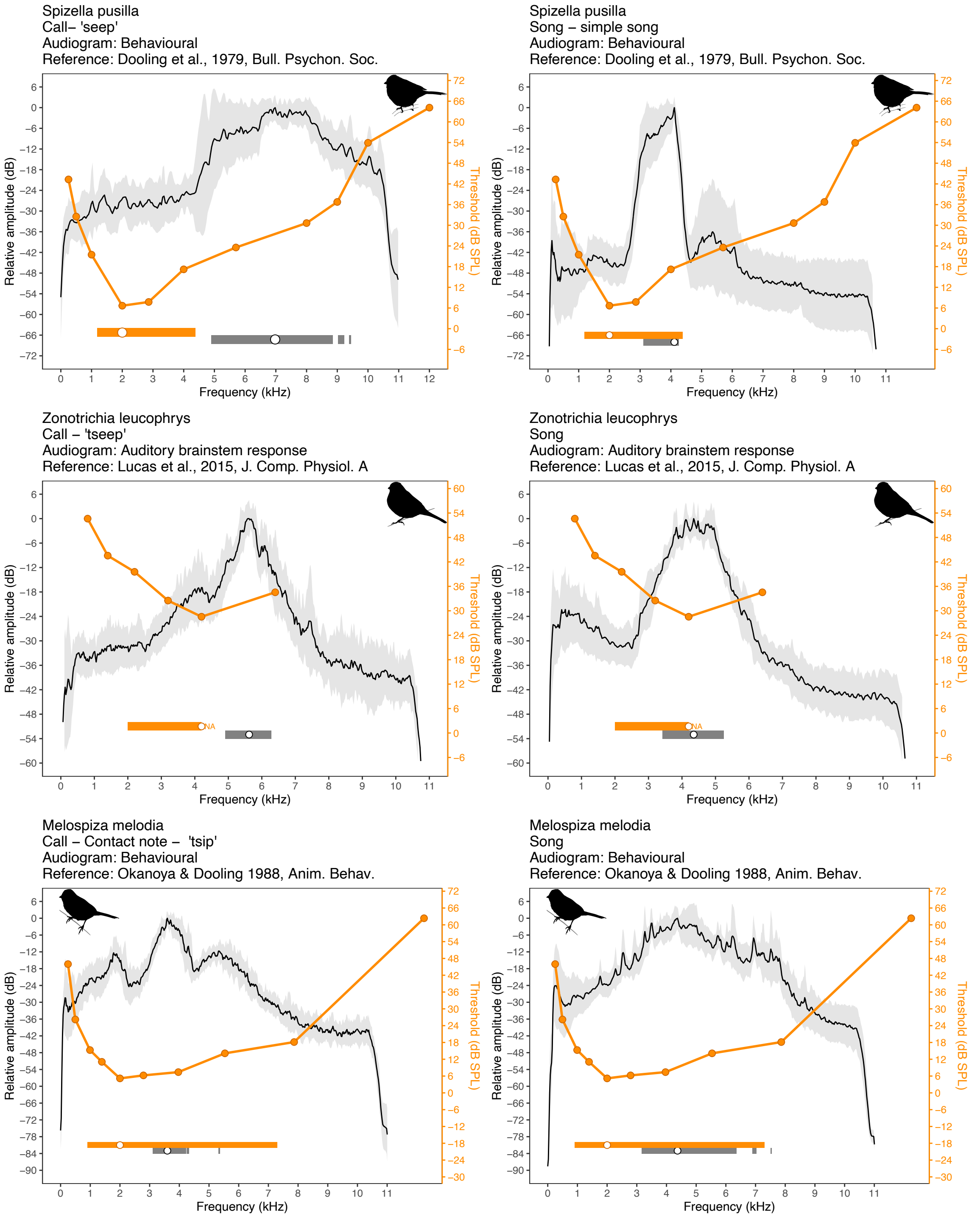


***(continued)*** **Fig. S15:** Audiograms (orange) and average power spectrum (black) for the order Passeriformes. They grey shading corresponds to the S.D. of the power spectra analyzed. Horizontal bars at the bottom show the auditory (orange) and vocal (grey) bandwidths (at 12 dB), and the points inside of them depict the best hearing frequency of the audiogram and the peak frequency of the mean spectrum.

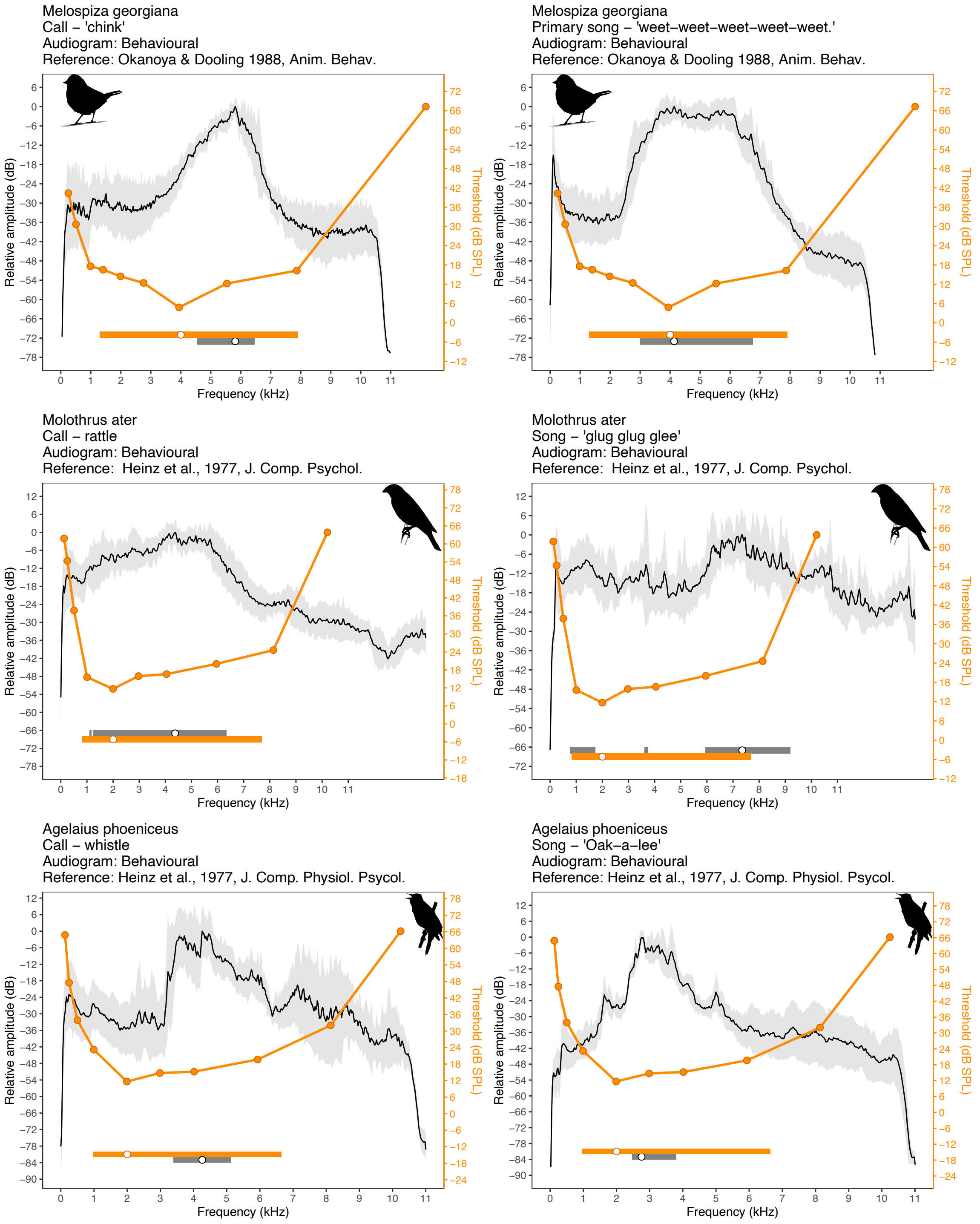


***(continued)*** **Fig. S15:** Audiograms (orange) and average power spectrum (black) for the order Passeriformes. They grey shading corresponds to the S.D. of the power spectra analyzed. Horizontal bars at the bottom show the auditory (orange) and vocal (grey) bandwidths (at 12 dB), and the points inside of them depict the best hearing frequency of the audiogram and the peak frequency of the mean spectrum.

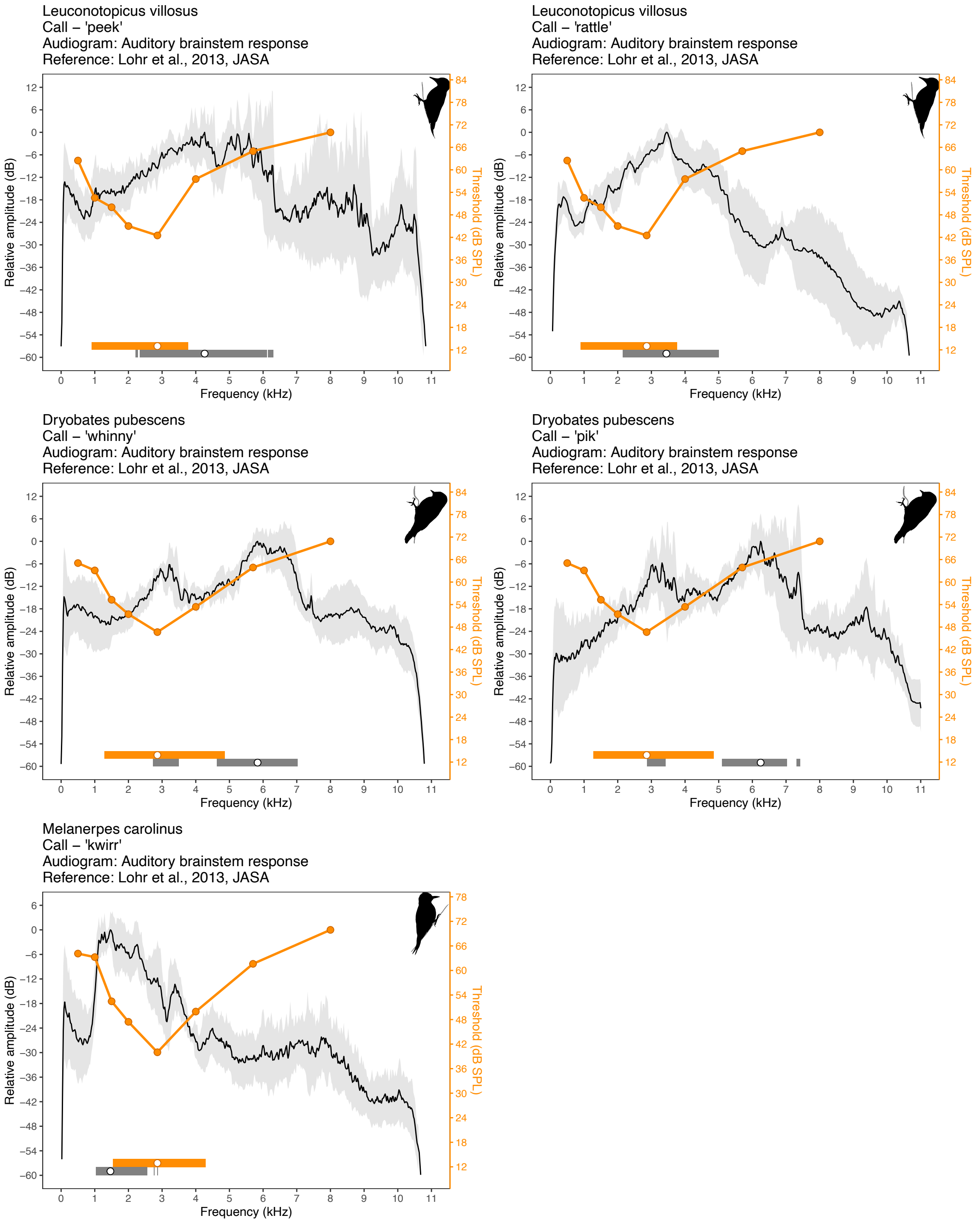


**Fig. S16:** Audiograms (orange) and average power spectrum (black) for the order Piciformes. They grey shading corresponds to the S.D. of the power spectra analyzed. Horizontal bars at the bottom show the auditory (orange) and vocal (grey) bandwidths (at 12 dB), and the points inside of them depict the best hearing frequency of the audiogram and the peak frequency of the mean spectrum.

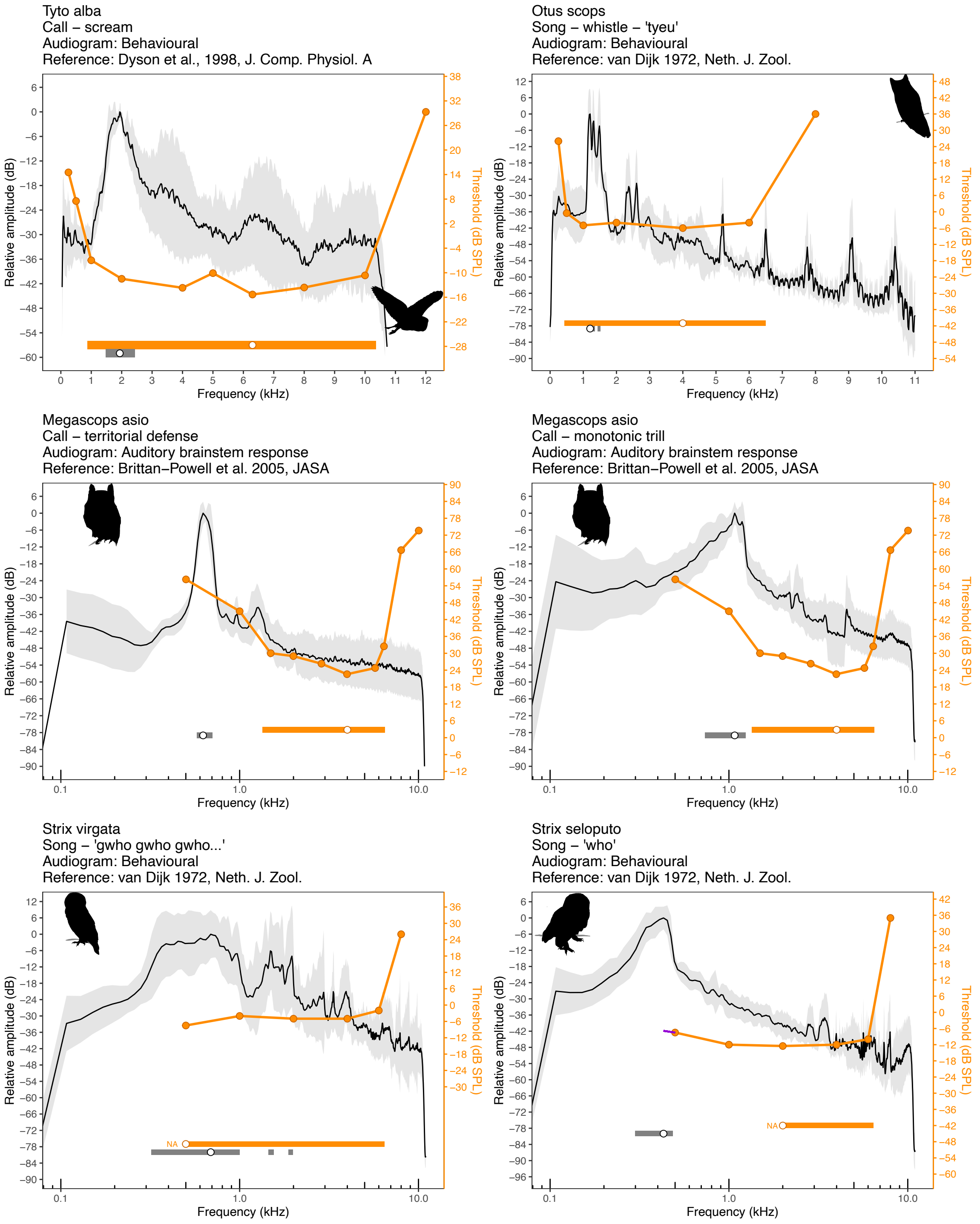


**Fig. S17:** Audiograms (orange) and average power spectrum (black) for the order Strigiformes. They grey shading corresponds to the S.D. of the power spectra analyzed. Horizontal bars at the bottom show the auditory (orange) and vocal (grey) bandwidths (at 12 dB), and the points inside of them depict the best hearing frequency of the audiogram and the peak frequency of the mean spectrum. The purple line in the audiogram of *Strix seloputo* shows the linear extrapolation used to estimate the threshold at the peak frequency of their song.

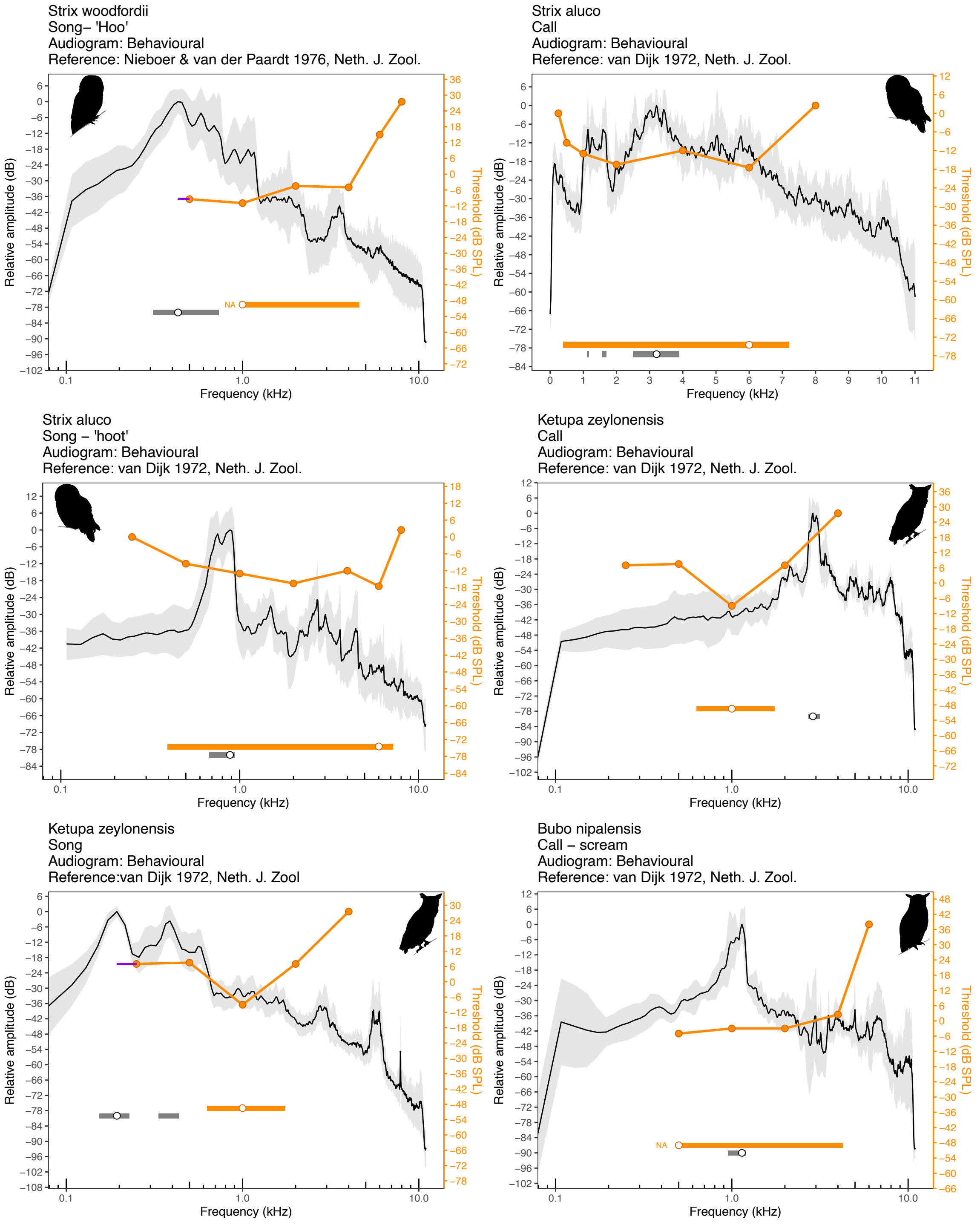


***(continued)*** **Fig, S17:** Audiograms (orange) and average power spectrum (black) for the order Strigiformes. They grey shading corresponds to the S.D. of the power spectra analyzed. Horizontal bars at the bottom show the auditory (orange) and vocal (grey) bandwidths (at 12 dB), and the points inside of them depict the best hearing frequency of the audiogram and the peak frequency of the mean spectrum. The purple line in the audiogram of *Strix woodfordii* and *Ketupa zeylonensis* shows the linear extrapolation used to estimate the threshold at the peak frequency of their songs.

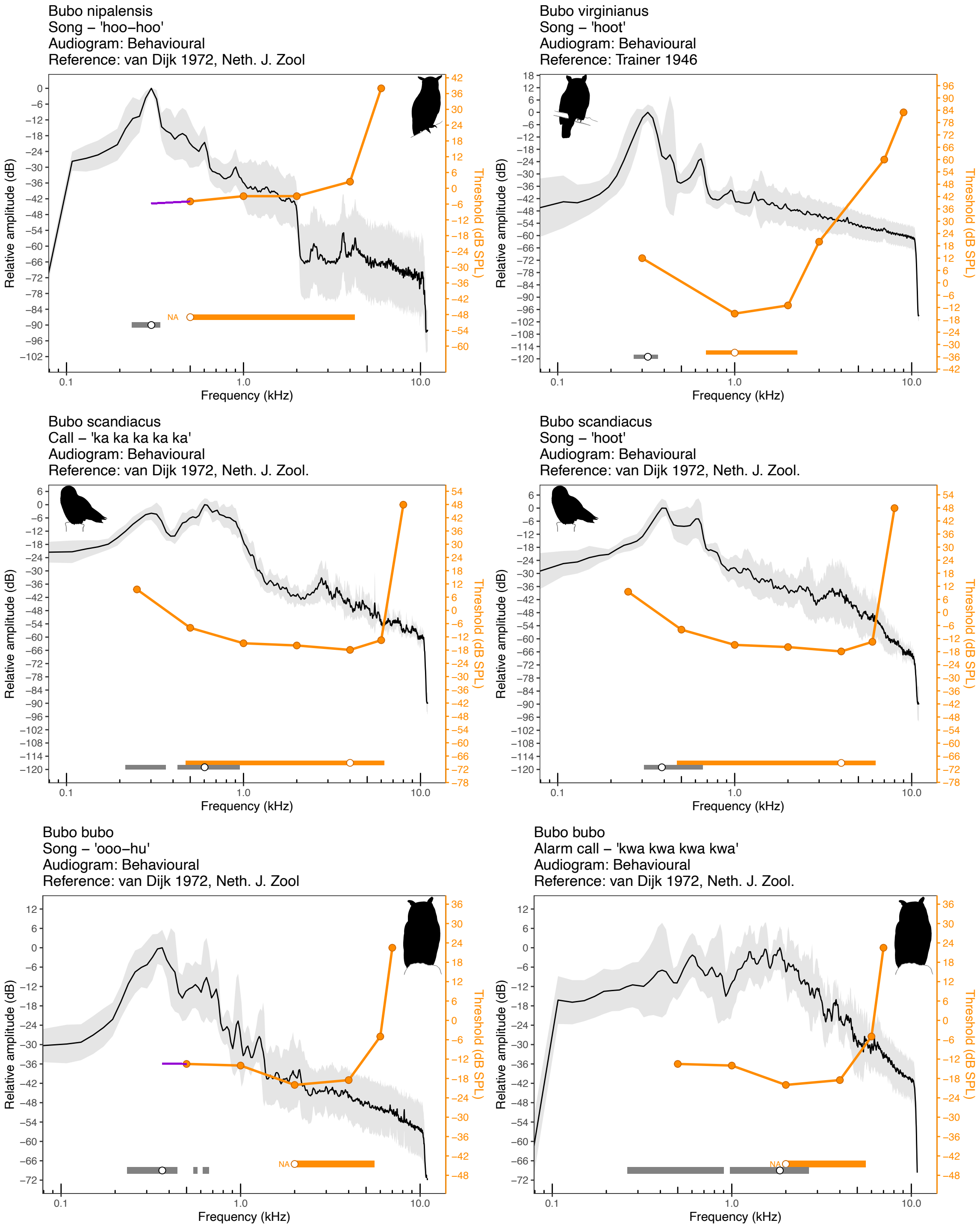


***(continued)*** **Fig. S17:** Audiograms (orange) and average power spectrum (black) for the order Strigiformes. They grey shading corresponds to the S.D. of the power spectra analyzed. Horizontal bars at the bottom show the auditory (orange) and vocal (grey) bandwidths (at 12 dB), and the points inside of them depict the best hearing frequency of the audiogram and the peak frequency of the mean spectrum. The purple line in the audiogram of *Bubo nipalensis* and *Bubo bubo* shows the linear extrapolation used to estimate the threshold at the peak frequency of their songs.

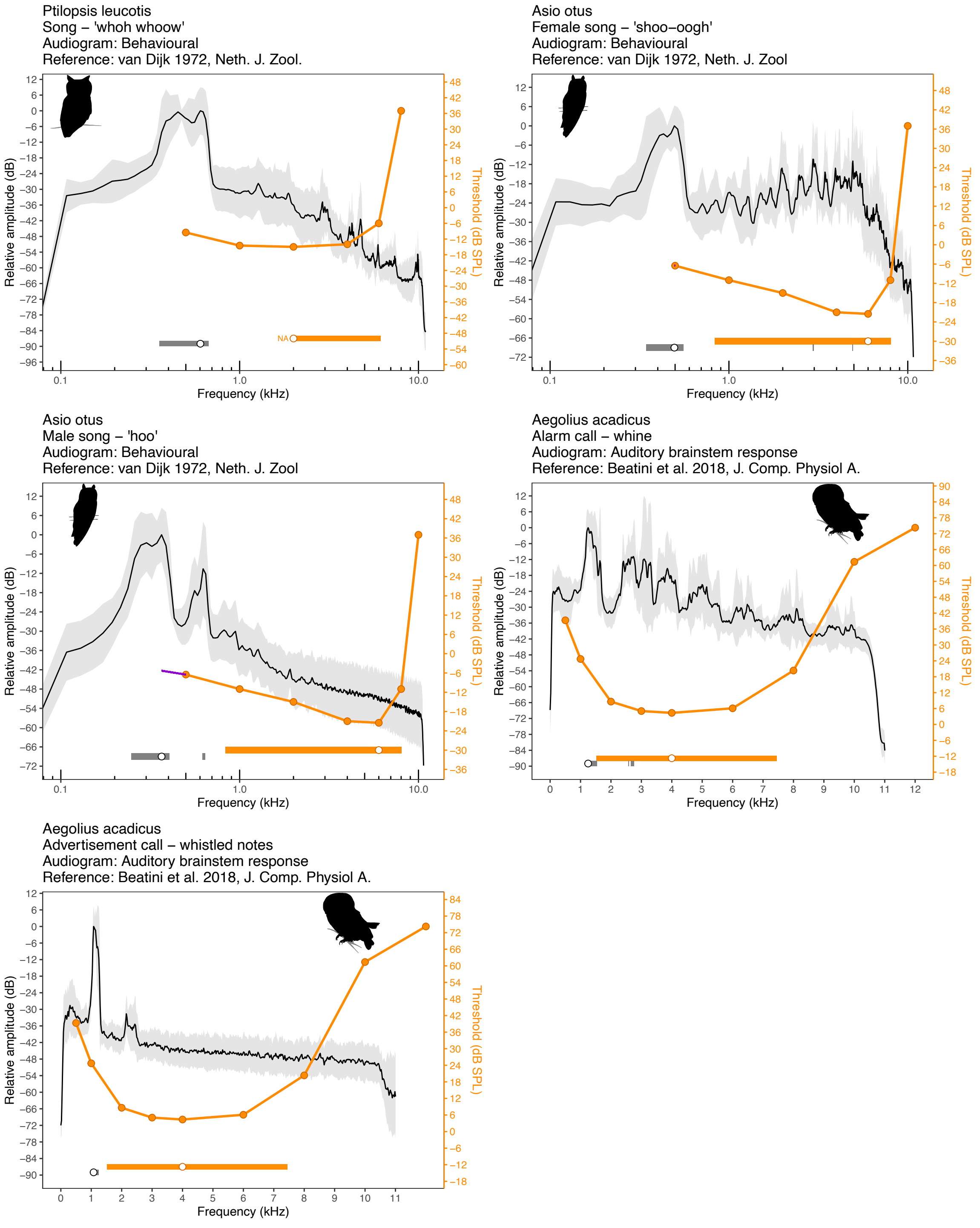


***(continued)*** **Fig. S17:** Audiograms (orange) and average power spectrum (black) for the order Strigiformes. They grey shading corresponds to the S.D. of the power spectra analyzed. Horizontal bars at the bottom show the auditory (orange) and vocal (grey) bandwidths (at 12 dB), and the points inside of them depict the best hearing frequency of the audiogram and the peak frequency of the mean spectrum. The purple line in the audiogram of *Asio otus* shows the linear extrapolation used to estimate the threshold at the peak frequency of the male song.

Haliaeetus leucocephalus
Scream – 'peal call'
Audiogram: Auditory brainstem response
Reference: McGee et al. 2019, J. Comp. Physiol. A

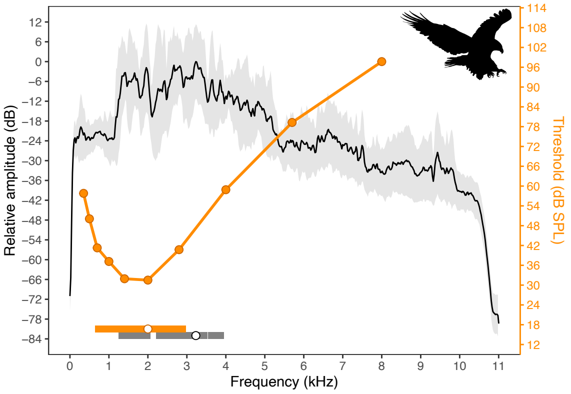


Haliaeetus leucocephalus
Keek – chatter call – 'kwit kwit kwit kwit kee–kee–kee–kee–ker'
Audiogram: Auditory brainstem response
Reference: McGee et al. 2019, J. Comp. Physiol. A

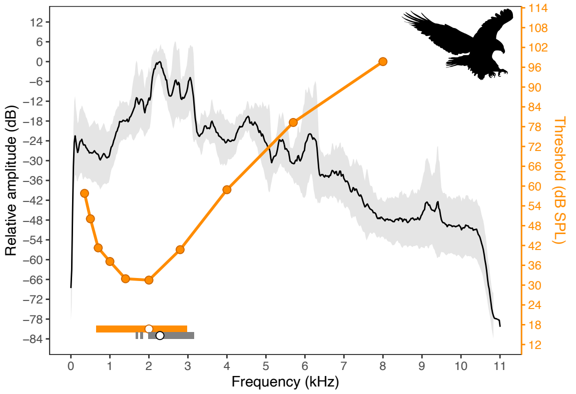


Buteo jamaicensis
Call – scream – 'kee–eeee–arrr'
Audiogram: Auditory brainstem response
Reference: McGee et al. 2019, J. Comp. Physiol. A

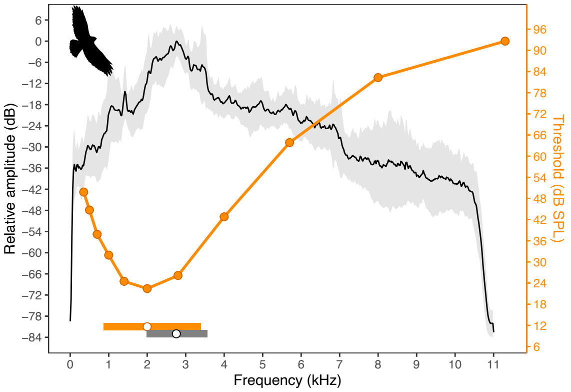


Accipiter nisus
Call – 'scream'
Audiogram: Behavioural
Reference: Klump et al., 1986, Behav. Ecol. Sociobiol.

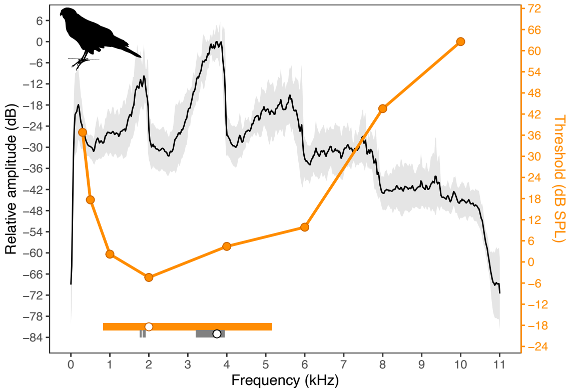


Accipiter nisus
Alarm call – 'kek kek kek kek kek'
Audiogram: Behavioural
Reference: Klump et al., 1986, Behav. Ecol. Sociobiol.

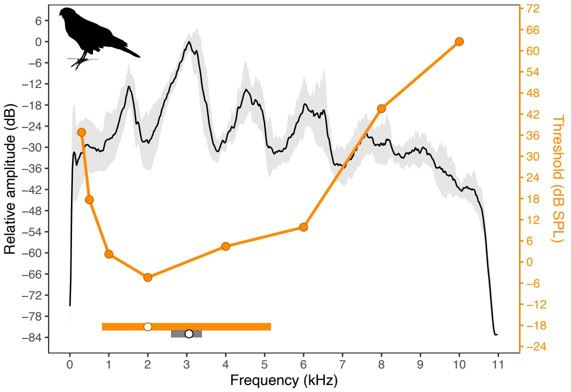


**Fig. S18:** Audiograms (orange) and average power spectrum (black) for the order Accipitriformes. They grey shading corresponds to the S.D. of the power spectra analyzed. Horizontal bars at the bottom show the auditory (orange) and vocal (grey) bandwidths (at 12 dB), and the points inside of them depict the best hearing frequency of the audiogram and the peak frequency of the mean spectrum.

**Table S1.** Species included in the study and methods used to estimate their audiograms. ABR = auditory brainstem response.

| Order | Common name | Scientific name | Reference | Method |
|---|---|---|---|---|
| Accipitriformes | Eurasian sparrowhawk | *Accipiter nisus* | (1) | Behavioral |
| Accipitriformes | Bald eagle | *Haliaeetus leucocephalus* | (2) | ABR |
| Accipitriformes | Red-tailed hawk | *Buteo jamaicensis* | (2) | ABR |
| Anseriformes | Long-tailed ducks | *Clangula hyemalis* | (3) | ABR |
| Anseriformes | White-winged scoters | *Melanitta fusca* | (3) | ABR |
| Anseriformes | Black scoters | *Melanitta americana* | (3) | ABR |
| Anseriformes | Harlequin ducks | *Histrionicus histrionicus* | (3) | ABR |
| Anseriformes | Ruddy ducks | *Oxyura jamaicensis* | (3) | ABR |
| Anseriformes | Common eiders | *Somateria mollisima* | (3) | ABR |
| Anseriformes | Lesser scaup | *Aythya affinis* | (4) | Behavioral + ABR |
| Anseriformes | Mallard duck | *Anas platyrhynchos* | (5) | Behavioral |
| Apodiformes | Blue-throated hummingbird | *Lampornis clemenciae* | (6) | ABR |
| Charadriiformes | Puffin | *Fratercula arctica* | (7) | ABR |
| Charadriiformes | Common murre | *Uria aalge* | (8) | ABR |
| Charadriiformes | Murrelet | *Brachyramphus marmoratus* | (9) | ABR |
| Columbiformes | Pidgeon | *Columba livia* | (10) | Behavioral |
| Falconiformes | American kestrel | *Falco_spaverius* | (11) | Behavioral |
| Galliformes | Indian peafowl | *Pavo cristatus* | (12) | Behavioral |
| Galliformes | Helmeted guineafowl | *Numida meleagris* | (13) | Behavioral |
| Galliformes | Chicken | *Gallus gallus domesticus* | (14) | Behavioral |
| Galliformes | Japanese quail | *Coturnix japonica* | (15) | Behavioral |
| Galliformes | Bobwhite quail | *Colinus virginianus* | (16) | Behavioral |
| Galliformes | Turkey | *Meleagris gallopavo domesticus* | (17) | Behavioral |

| | | | | |
|---|---|---|---|---|
| Gaviiformes | Red-throated loons | *Gavia stellata* | (3) | ABR |
| Passeriformes | Chickadee | *Poecile carolinensis* | (18) | ABR |
| Passeriformes | Tufted titmice | *Baeolophus bicolor* | (19) | ABR |
| Passeriformes | House sparrows | *Passer domesticus* | (19) | ABR |
| Passeriformes | White-breasted nuthatches | *Sitta carolinensis* | (19) | ABR |
| Passeriformes | Common canary | *Serinus canarius* | (20) | Behavioral |
| Passeriformes | House finch | *Carpodacus mexicanus* | (21) | Behavioral |
| Passeriformes | Redwing blackbird | *Agelaius phoeniceus* | (22) | Behavioral |
| Passeriformes | Brown-headed cowbird | *Molothrus ater* | (22) | Behavioral |
| Passeriformes | Great tit | *Parus major* | (23) | Behavioral |
| Passeriformes | Red-billed firefinch | *Lagonosticta senegala* | (24) | Behavioral |
| Passeriformes | Swamp sparrow | *Melospiza georgiana* | (25) | Behavioral |
| Passeriformes | Song sparrow | *Melospiza melodia* | (25) | Behavioral |
| Passeriformes | Zebra finch | *Taeniopygia guttata* | (26) | Behavioral |
| Passeriformes | Blue jay | *Cyanocitta cristata* | (27) | Behavioral |
| Passeriformes | Field sparrow | *Spizella pusilla* | (28) | Behavioral |
| Passeriformes | Long-tailed finch | *Poephila acuticauda* | (29) | Behavioral |
| Passeriformes | Double-barred finch | *Taeniopygia bichenovii* | (29) | Behavioral |
| Passeriformes | Bengalese finch | *Lonchura striata domestica* | (29) | Behavioral |
| Passeriformes | Gold-breasted waxbill | *Amandava subflava* | (29) | Behavioral |
| Passeriformes | Hooded crow | *Corvus corone cornix* | (30) | Behavioral |
| Passeriformes | Bullfinch | *Pyrrhula pyrrhula* | (31) | Behavioral |
| Passeriformes | White crowned sparrow | *Zonotrichia leucophrys* | (32) | ABR |
| Piciformes | Downy woodpecker | *Picoides pubescens* | (33) | ABR |
| Piciformes | Hairy woodpecker | *Picoides villosus* | (33) | ABR |
| Piciformes | Red-bellied woodpecker | *Melanerpes carolinus* | (33) | ABR |

| | | | | |
|---|---|---|---|---|
| Psittaciformes | Cockatiel | *Nymphicus hollandicus* | (26) | Behavioral |
| Psittaciformes | Budgerigar | *Melopsittacus undulatus* | (34) | Behavioral |
| Psittaciformes | Kea | *Nestor notabilis* | (35) | Behavioral |
| Psittaciformes | Orange-fronted conure | *Aratinga canicularis* | (36) | Behavioral |
| Sphenisciformes | Backfooted penguin | *Spheniscus demersus* | (37) | Cochlear_potential |
| Steatornithiformes | Oilbird | *Steatornis caripensis* | (38) | Cochlear_potential |
| Strigiformes | Saw-whet owl | *Aegolius acadicus* | (39) | ABR |
| Strigiformes | Eastern Screech Owl | *Megascops asio* | (40) | ABR |
| Strigiformes | African woodowl | *Strix woodfordii* | (41) | Behavioral |
| Strigiformes | European bawn owl | *Tyto alba guttata* | (42) | Behavioral |
| Strigiformes | Tawny owl | *Strix aluco* | (43) | Behavioral |
| Strigiformes | Long-eared owl | *Asio otus* | (43) | Behavioral |
| Strigiformes | Common scops owl | *Otus scops* | (43) | Behavioral |
| Strigiformes | White-faced scops owl | *Otus leucotis* | (43) | Behavioral |
| Strigiformes | Eagle-owl | *Bubo bubo* | (43) | Behavioral |
| Strigiformes | Forest eagle-owl | *Bubo nipalensis* | (43) | Behavioral |
| Strigiformes | Brown fish owl | *Ketupa zeylonensis* | (43) | Behavioral |
| Strigiformes | Snowy owl | *Nyctea scandiaca* | (43) | Behavioral |
| Strigiformes | Mottled owl | *Strix virgata* | (43) | Behavioral |
| Strigiformes | Spotted wood owl | *Strix seloputo* | (43) | Behavioral |
| Strigiformes | Great horned owl | *Bubo_virginianus* | (11) | Behavioral |
| Suliformes | Northern gannet | *Morus bassanus* | (3) | ABR |
| Suliformes | Great cormorant | *Phalacrocorax carbo* | (44) | Behavioral |
